# Atypical plastid genome evolution: *Cereus* Mill. from distinct environments harbor one of the largest plastid genomes in Cactaceae

**DOI:** 10.1101/2025.08.18.670840

**Authors:** Clara C.V. Badia, Maria C. Silva, Valter A. de Baura, Eduardo Balsanelli, Emanuel Maltempi de Souza, Fábio de Oliveira Pedrosa, Jéferson Fregonezi, Marcelo Rogalski

**Affiliations:** Laboratório de Sistemática e Evolução de Plantas, Departamento de Biologia Vegetal, Universidade Federal de Viçosa, Viçosa-MG, Brasil; Laboratório de Fisiologia Molecular de Plantas, Departamento de Biologia Vegetal, Universidade Federal de Viçosa, Viçosa-MG, Brasil; Núcleo de Fixação Biológica de Nitrogênio, Departamento de Bioquímica e Biologia Molecular, Universidade Federal do Paraná, Curitiba-PR, Brasil

**Keywords:** Tribe Cereeae, Plastid evolution, Molecular markers, RNA editing

## Abstract

**Background:** Cactaceae has successfully radiated in xeric habitats across the Americas, presenting very distinct morphologies and evolutionary patterns within tribes. This study presents the complete plastomes of *C. jamacaru* subsp. *jamacaru* and *C. hildmannianus* subsp. *hildmannianus*, which inhabit distinct habitats, providing insights into their genomic structure and evolutionary history, with implications for conservation.

**Methods and Results:** Chloroplast genomes of the two *Cereus* were assembled and analyzed to investigate plastome evolution in Cactoideae. Fresh cladodes were collected and their mesophyll manually extracted, chloroplasts were extracted from the mesophyll, and cpDNA sequenced using *Illumina MiSeq*. *De novo* assembly and annotation were conducted using BLAST, Expasy, and tRNAScan as validation tools. We compared the genome structure, gene content, codon usage, and RNA editing predictions between tribes. The genome was 141.884 and 141.600 bp for *C. jamacaru* and *C. hildmannianus,* respectively, and dotplot analysis confirmed highly syntenic plastomes. The genes *trnV-GAC, trnV-UAC, rpl23, ndhA, ndhE, ndhG, ndhI,* and *ndhK* were lost, and *ndhB, ndhC, ndhF,* and *rpl33* are pseudogenes. The tRNA^val^ losses indicate putative superwobbling or nuclear-coded tRNA import from cytosol. We identified an insertion in *rps18* for both *Cereus*, suggesting that intron retention may be in course for these species. We identified ∼190 single sequence repeats and 50 tandem repeats for each species, and eight exclusive RNA editing sites. Synteny analysis revealed rearrangements distinguishing taxa within Cactoideae. Phylogenetic results supported *Cereus* monophyly, corroborating existing classifications, and clarifies unresolved relationships, enhancing understanding of phylogenetic relationships within Cactaceae.

**Conclusions:** Our results provide evidence on the evolutionary patterns and putative signatures of adaptation to distinct environments, providing insights into genomic evolution and conservation of *Cereus*.

## Introduction

Cactaceae stands out as an informative model for understanding the diversification in xeric habitats due to its successful radiation in the New World [1]. Estimated to have arisen around 35 million years ago (Ma), with most speciose clades originating more recently, around 10-5 Ma [1, 2], the family has about 1,450 species and 129 genera [3]. The occurrence of the species is concentrated in the Americas, with the main centers of diversity being Mexico, the Andes, and the northeastern region of Brazil [4].

Although it is challenging to determinate the key drivers of such successful radiation, the accentuated adaptations of cacti to the most distinct environments, including deep modifications in lifestyles over the course of evolution, may explain the wide distribution of Cactaceae representatives [2, 5, 6]. Despite having a wide distribution, Cactaceae is among the five most endangered families, with more than 30% of its species threatened [7].

Cactoideae is the most species-rich subfamily of Cactaceae and is subdivided into 9 tribes that encompass a vast range of lifestyles and growth forms (e.g., treelike, shrubby, caespitose, climbing, or epiphytes) [2]. It has relatively high convergence rates of growth forms within its tribes, as is the case of the “BCT clade”, which refers to Browningieae, Cereeae, and Trichocereeae [4]. This clade spans a broad range of growth-forms, being barrel, erect and ribbed the main ones. Each one of these is differently inherited throughout this larger clade, therefore presenting distinct representativeness across the *taxa*.

Cereeae comprises the subject of this study, *Cereus* Mill., which has Brazil as its main center of diversity. The genus comprises 18 species, 8 being endemic to Brazil; and 6 subspecies. Reconstructions revealed that *Cereus* initially diversified from the Cerrado towards the Caatinga around 3.5 Ma, and that the group comprising the species of interest of this paper have diversified just over 2 million years ago, making it the most recently radiated [8].

*Cereus jamacaru* DC., popularly known as “Mandacaru”, is typical of the semiarid region comprised in the northeastern region of Brazil (**Figure 1**). This species is of utmost importance in terms of characterizing the Caatinga phytogeographical domain, local culture and economy [1, 7]. Mandacaru is characterized by its arboreal habit and rapid growth and is commonly found in various physiognomies of the Caatinga, usually associated with rock outcrops [9]. *Cereus hildmannianus* K.Schum. subsp. *hildmannianus*, “Tuna”, occurs in the south-east and south of the Neotropics, specifically in the Pampa, a xeric environment where low temperatures are commonly recorded (**Figure 2**). It is often found in semi-deciduous ombrophilous forests associated with rock outcrops [9]. Tuna occasionally occurs as an epiphyte on trees and shrubs as well [10].

**Figure 1.**
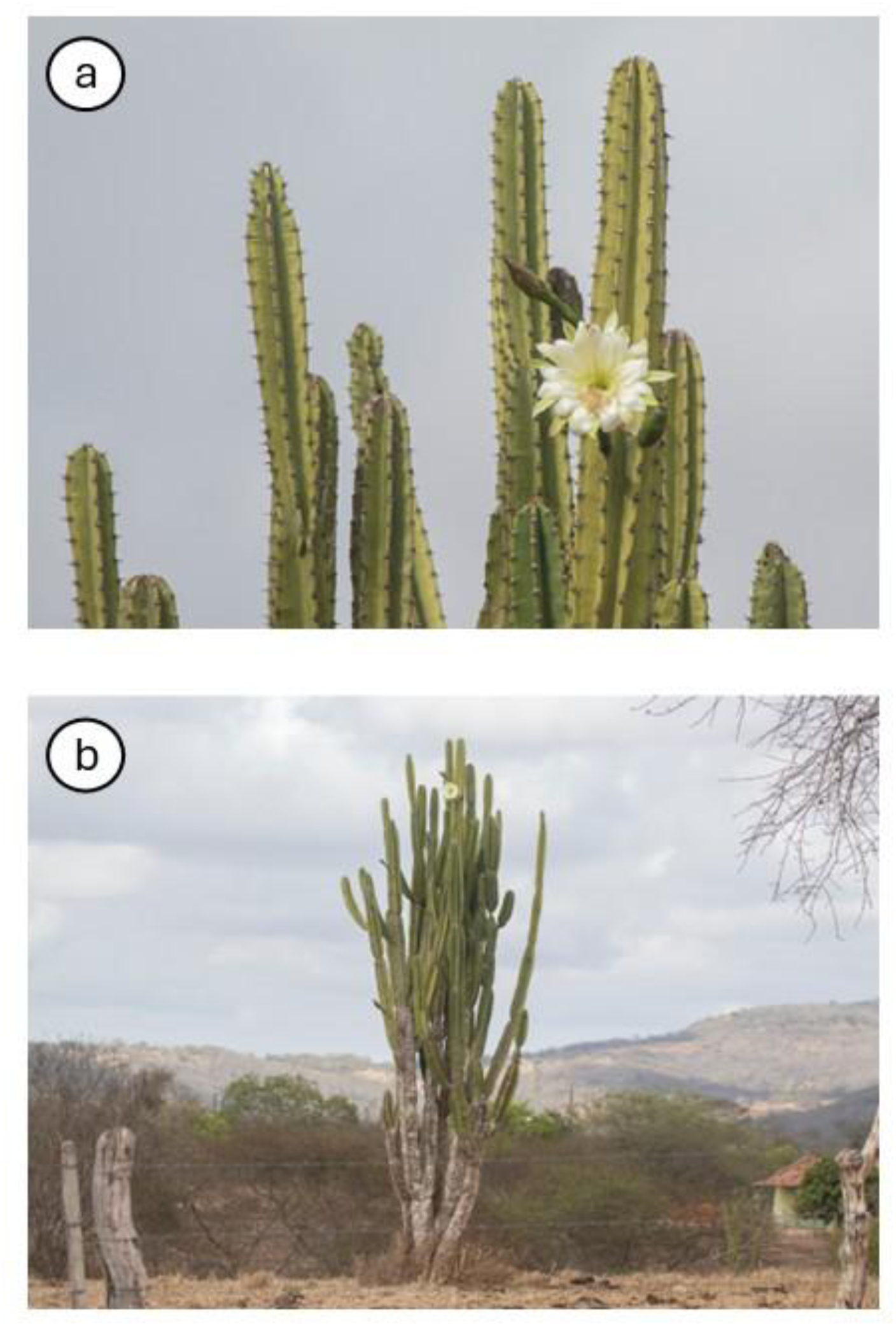
Morphology (a) and habit (b) of Ce*reus jamacaru* subsp. *jamacaru*. Photos: Alenilson Rodrigues.

**Figure 2.**
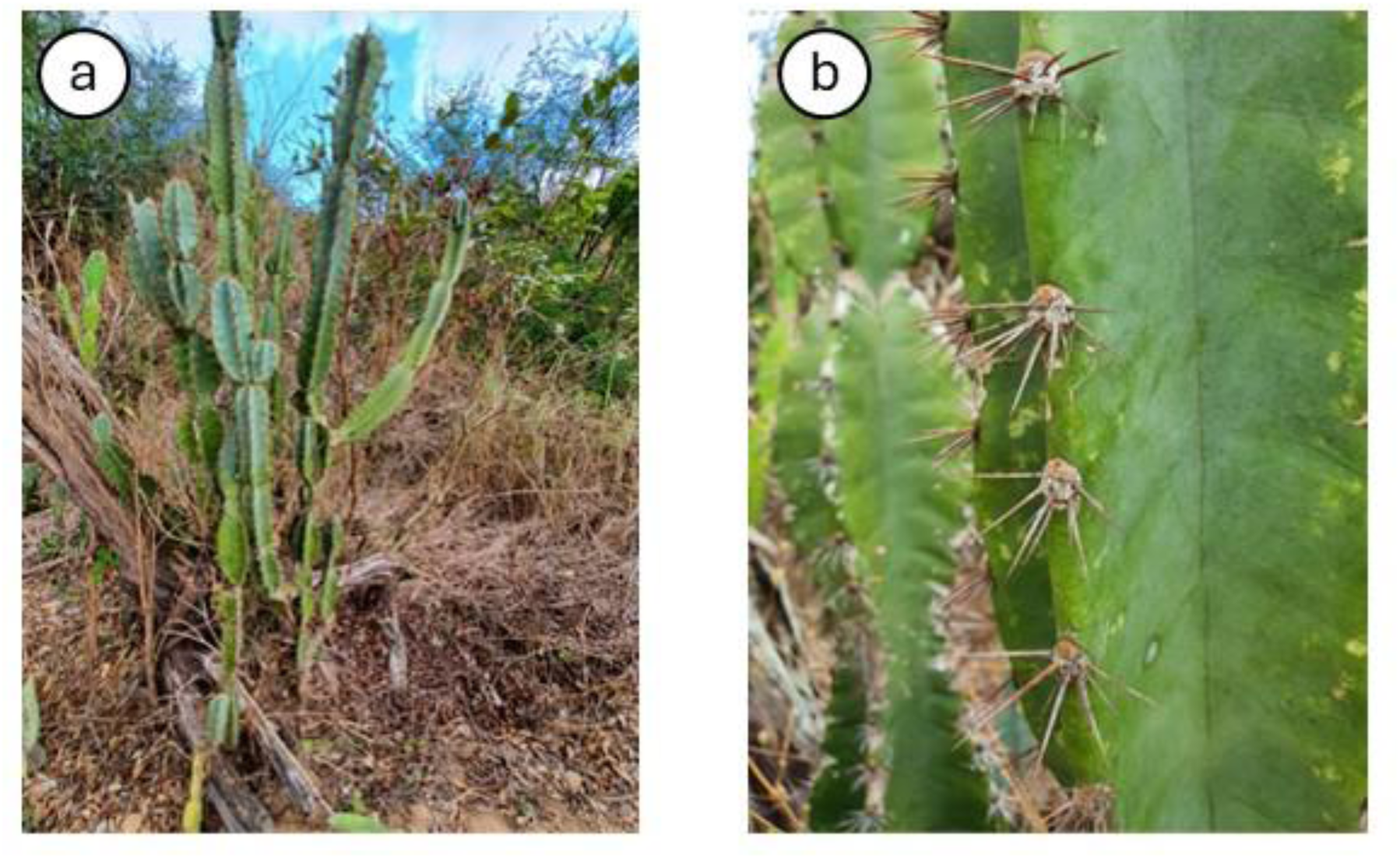
Habit (a) and morphology (b) of Ce*reus hildmannianus* subsp. *hildmannianus*. Photos: Caio Crelier.

Recent investigations into the genus *Cereus* have employed various molecular approaches, including single sequence repeats (SSRs) transferability, phylogenetic inferences [11], and phylogeographic structure [12]. In this context, the whole plastid genome provides broader evidence when compared to intergenic spacers and/or non-coding regions previously used for molecular studies in the genus. Growing knowledge of these species is especially interesting when dealing with species whose conservation status is unknown, despite their economic and ecological importance.

Herein we present the complete plastome of *Cereus jamacaru* (hereafter: *C. jamacaru*) and *Cereus hildmannianus* subsp. *hildmannianus* (hereafter: *C. hildmannianus*), detailing its structure and genetic content and order, and placing their genomic characteristics within the subfamily in a comparative evolutionary frame. We compared both genomes framing them in a contrasting habitat scenario, since they inhabit strikingly different environments.

## Materials and methods

### Plant material, chloroplast isolation, and plastid DNA extraction

Fresh and young cladodes of *Cereus jamacaru* and *C. hildmannianus* were collected from adult specimens grown in a greenhouse and their mesophyll prepared for extraction as previously outlined [14]. Prior to chloroplast isolation, the cladodes were stored at 4°C in the dark for five days to reduce the starch content. Chloroplast isolation was conducted first, and cpDNA extraction were then performed following the methods available in the literature [14]. The cpDNA was sent to the Federal University of Paraná (UFPR), where it was sequenced according to the following section.

### Sequencing, assembly, annotation, and data archiving statement

Around 1 ng of plastidial DNA was used to prepare sequencing libraries using the Nextera XT DNA Sample Prep Kit (Illumina Inc., San Diego, CA, USA) according to the manufacturer’s guidelines. Libraries were sequenced on an Illumina MiSeq platform (Illumina Inc., San Diego, CA, USA) using a paired-end approach with 2 x 250 bp read configuration.

The sequencing of three technical replicates of *C. jamacaru* obtained a total of 57.582 reads ranging from 22 to 41.776 bp in five contigs. The average coverage of contigs used for assemblage ranged from 31.23 to 394.33, presenting an overall average of 248.09. The reads were trimmed under the threshold with >0.05 of probability of error. Subsequently, the trimmed reads were *de novo* assembled in contigs in CLC genomics Workbench 8.0.2 (CLC Bio, Aarhus, Denmark) and the resulted contigs were manually curated.

The sequencing of two technical replicates of *C. hildmannianus* obtained 46.336 reads ranging from 19 to 32.334 bp. We performed *de novo* assembly of the trimmed reads in contigs in CLC Genomics Workbench 8.0.2 software (CLC Bio, Aarhus, Denmark). We used initially three contigs to assemble the plastome of *Cereus hildmannianus*, ranging from 361,45 to 3.221,27 of average coverage, presenting an overall average of 2.148,93. Two gaps were identified using the contigs generated by CLC Genomics, of 258 bp and 387 bp, so we performed a second round of *de novo* assembly in NOVOPlasty v. 4.2 [15] for both replicates. It resulted in 1.592.944 reads and five contigs ranging from 10.319 to 30.393 bp for the first replicate; and 1.492.984 reads and 7 contigs ranging from 2.522 to 30.084 bp for the second one. The gap of 258 bp was successfully closed using a contig of 30.393 bp and average coverage of 48.637, whereas the gap of 387 bp was solved by a contig of 30.083 bp and average coverage of 21.535. The closure of the latter gap was additionally supported by a second contig with same size and gene content and order, and average coverage of 116.118.

For the assembly of both *Cereus,* conflicting nucleotides among overlapping or homologous contigs were inspected and resolved to ensure sequence accuracy. The average coverage per nucleotide was meticulously checked in CLC genomics Workbench 8.0.2 software to keep the one presenting the highest quality score. The boundaries of the Inverted Repeats (IRA and IRB) were identified using REPuter, with parameters set to a hamming distance of 3 and minimum repeat size ranging from 30 to 100 bp.

We used the Annotation of Organellar Genomes (GeSeq) for an initial annotation of the genes [16]. The annotations were manually checked using the BLAST tool available in NCBI and subsequently compared to related species via MAFFT alignment to check for initiation, termination, and intron codon positions. Gene losses identified during the annotation process were checked manually by searching regions throughout the plastome that were similar to those genes in the reference species *Spinacia oleracea* (NC_002202), and *Portulaca oleracea* (NC_041264). We used tRNAscan-SE webserver with search mode set to default [17] to check for tRNA annotation and respective functionality. The plastome map was drawn in the webserver Organellar Genome DRAW (OGDRAW) [18]. The complete nucleotide sequences of *C. jamacaru* and *C. hildmannianus* plastomes are available in the GenBank database of NCBI under the accessions PV032414 and PQ997910 respectively.

### Structural characterization of the plastomes

The structural features of the *Cereus jamacaru* and *Cereus hildmannianus* plastomes were analyzed in MAUVE progressive alignment of whole genomes (Darling et al. 2004) embedded in Geneious v. 2022.1. In the absence of a sequence in GenBank for the closest species available in the database under the “verified” status (*Cereus fernambucensis,* OL397055.1), we compared both plastomes to the *Melocactus glaucescens’* (OK298499, also belonging to the tribe Cereeae) and to the Large Single Copy region of *Portulaca oleracea* (NC_041264) as reference, which spans from the *trnH-GUG* to the *rps19*. The linear plastome maps were generated in OGDRAW [18]. Synteny between the plastomes of the *Cereus* species and *Melocactus glaucescens* was evaluated in NUCmer (NUCleotide MUMmer) 3.23 [19].

### Codon Usage

Codon usage bias was analyzed for the protein-coding gene set of both *Cereus* species using the Codon Usage tool within the Sequence Manipulation Suite webserver (available at https://www.bioinformatics.org/sms2/codon_usage.html), applying the standard genetic code. The codon usage data of *C: jamacaru* and *C. hildmannianus* was then compared to members of the tribe Cereeae (namely *Cereus fernambucensis, Melocactus glaucescens*, and *Melocactus ernestii*). Additionally, to obtain a comprehensive overview of codon frequencies within the subfamily, we calculated the average and standard deviation for seven Cactoideae tribes and the Opuntioideae. Furthermore, we examined the raw codon usage fractions for *Portulaca oleracea* and *Spinacia oleracea*, which served as reference points (see **Supplementary Table S1** for the included accessions).

### Prediction of RNA editing sites

We accessed putative RNA editing sited in protein-coding genes of *C. hildmannianus* and *C. jamacaru* using the Predictive RNA Editor for Plants (PREP) [20]. Twenty-seven genes were analyzed, considering the availability of the webserver and excluding from this analysis the highly divergent genes such as *accD* and *clpP,* and absent/pseudogenes. The genes included were: *atpA, atpB, atpF, atpI, ccsA, matK. petB, petD, petG, petL, psaB, psaI, psbB, psbE, psbF, psbL, rpl2, rpl20, rpoA, rpoB, rpoC1, rpoC2, rps2, rps8, rps14, rps16,* and *ycf3*). The cut-off value was set to 0.8 for this predictive analysis.

### Simple Sequence Repeats (SSRs) and Tandem Repeats

We mapped the Simple Sequence Repeats (SSRs) in MIcroSAtellite (MISA) Identification webserver (https://webblast.ipk-gatersleben.de/misa/). The settings were as follows: eight repeat units for mononucleotide SSRs, four for di- and trinucleotide SSRs, and three for tetra-, penta-, and hexanucleotide SSRs. The maximum length of sequence between two SSRs to register as compound SSR was set to 0. We used Phobos Tandem Repeat Finder v. 1.0.6 embedded in Geneious v 2022.1. to infer the number, size and position of tandem repeats. The parameters of minimum and maximum repeat unit length were set to 1 and 200 respectively. To infer the size of the inverted repeats we used the webserver REPuter [21], in which minimal repeat size was set to 30 bp, identity of repeats ≥ 90% and hamming distance = 3. After mapping the SSRs and Tandem Repeats, we checked the location in the plastome to match the corresponding positions in Geneious v. 2022.1. In both analysis the Inverted Repeat A was removed to avoid redundancy.

### Phylogenetic inference

We reconstructed the phylogenetic relationships for the two species of *Cereus* and additional 56 *taxa*, including species belonging to the Cactineae suborder, representatives of the families Basellaceae, Montiaceae, Halophytaceae, Portulacaceae, and Talinaceae. We used *Spinacia oleracea* (Caryophyllales, same order as Cactaceae) as an outgroup, manually rooting the tree at this terminal after running the tree. The complete list of species and respective GenBank accessions included are available in **Supplementary Table S2**. We carefully checked for any misannotated genes within the sampled plastomes using the same pipeline as the one we used for the annotation of the *Cereus* plastomes (see the sequencing and annotation section for detail). After gathering correct information of the 45 protein-coding genes, we performed gene-by-gene alignment using MAFFT embedded in Geneious. After that, the concatenation of the gene alignments was carried out in Geneious, prior to the best-fit-model estimation for the partitions. A customized gap-removal script was developed in R v. 4.3.3 during data treatment using the packages *ape* and *seqinr* [22, 23].

In IQTree v.2.3.2 we first detected the partition models [24] allowing the partitions to have their own evolution rate and share the same set of branch lengths. The best substitution models and genes belonging to each of the 6 partitions identified by ModelFinder2 embedded in IQTree v.2.3.2 were, respectively: TVM+F+I+R2: *atpA, atpB, atpF, petA, psaC, psbH, psbT, rbcL, rps4*, and *ycf3*; TVM+F+I+R2: *atpE, ccsA, matK, psbJ, rpl14, rpl2, rpoA, rpoC2, rps12, rps14, rps2*, and *rps8*; TVM+F+I+G4: *atpH, atpI, petB, petD, petG, psaA, psaB, psaI, psbC, psbD, psbE, psbF, psbN*, and *psbZ*; TPM3u+F+G4*: infA* and *rps19*; JC: *petN*; TVM+F+I+R2: *psbL, psbM*, *rpl16, rps11, rps15*, and *rps7*.

The Maximum Likelihood reconstruction was carried out having the parameters set to “-p” and with five hundred non-parametric bootstrap replications to evaluate branch supports. The consensus tree was visualized on FigTree v.1.4.4 software (http://tree.bio.ed.ac.uk/software/figtree/).

## Results

### General features of the plastomes of *Cereus jamacaru* and *C. hildmannianus* subsp. *hildmannianus*

The plastomes of *Cereus* present the typical quadripartite structure, containing two single copies (Large Single Copy and Small Single Copy – LSC and SSC respectively) interspersed by palindromic repeats (Inverted Repeat A and B – IRA and IRB). The plastome of *Cereus jamacaru* presents 141.884 bp and is 284 bp larger than the plastome of *C. hildmannianus* (141.600 bp), and both plastomes present identical gene content and order (**Figure 3**; **Tables 1, 2**). The IRs were the largest among all Cactaceae, even when compared to the largest genome in the family (*Opuntia quimilo*, 150.347 bp; **Table 3**). *Cereus jamacaru* presented larger LSC (53.632 bp) and IR (32.237 bp) sizes than *C. hildmannianus* (LSC: 53.310 bp; IR: 31.955 bp), which, in turn, presented larger SSC (24.320 bp, against 23.778 bp in *C. jamacaru*).

**Figure 3.**
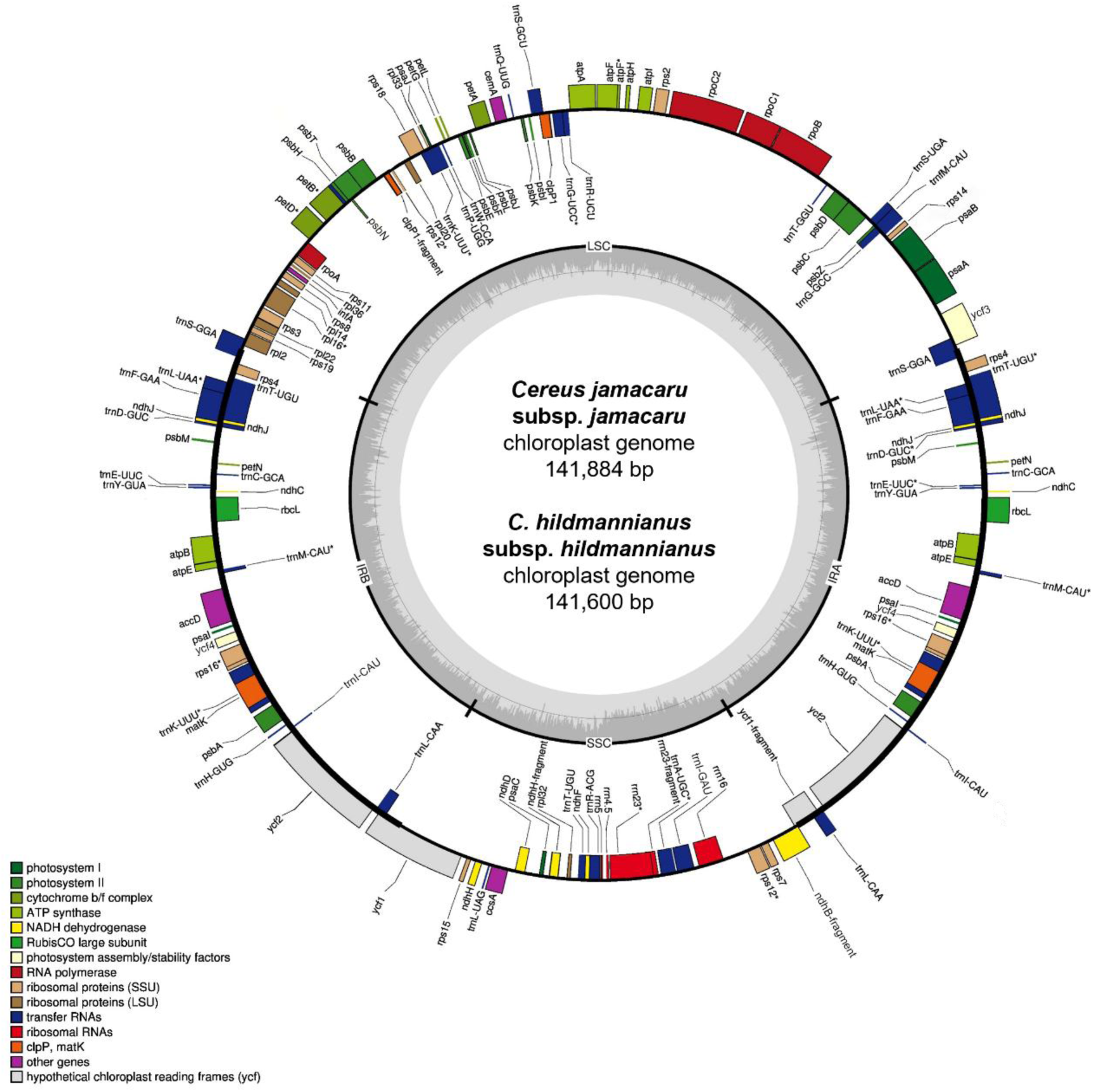
Gene map of *Cereus jamacaru* subsp. *jamacaru* and *C. hildmannianus* subsp. *hildmannianus* plastid genome. The circular DNA molecule is divided into four regions: a large single copy (LSC) region, a small single copy (SSC) region, and two inverted repeats (IR_A_ and IR_B_). Genes located in the inner side of the circle are transcribed in a clockwise direction, while those on the outer side are transcribed counterclockwise. The legend box on the left classifies genes into functional categories. Within the inner ring, darker gray indicates GC-rich regions, while lighter gray represents AT-rich regions. The dashed circle between the two shades denotes the 50% GC/AT content threshold.

**Table 1.**
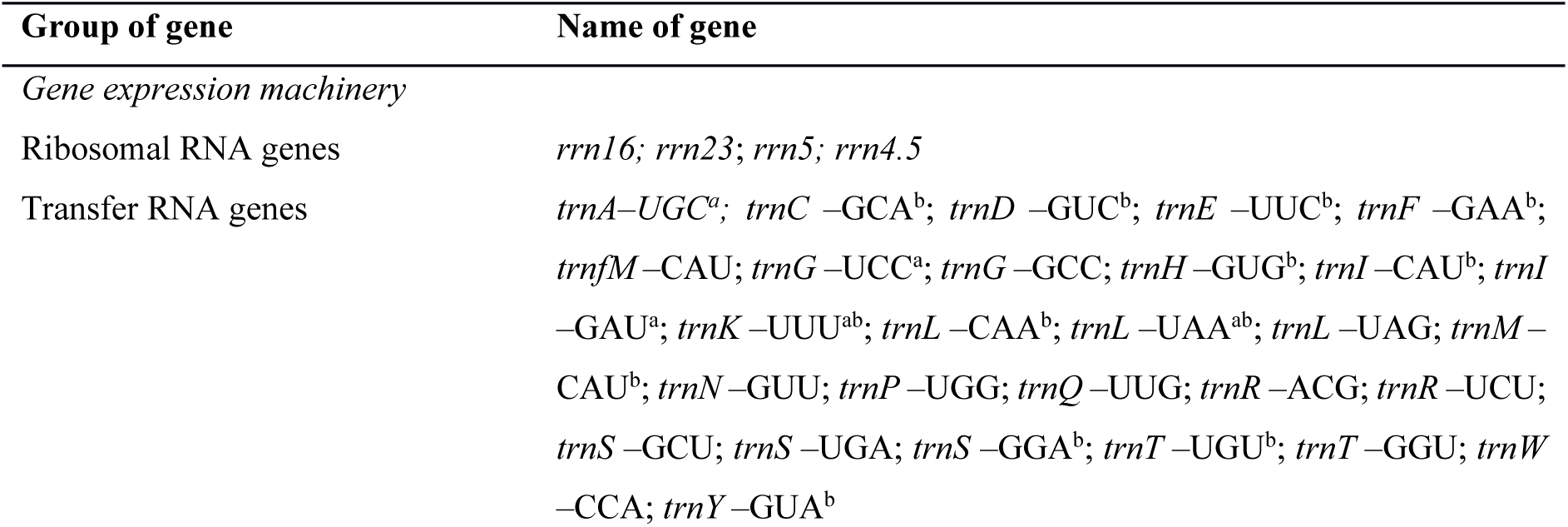
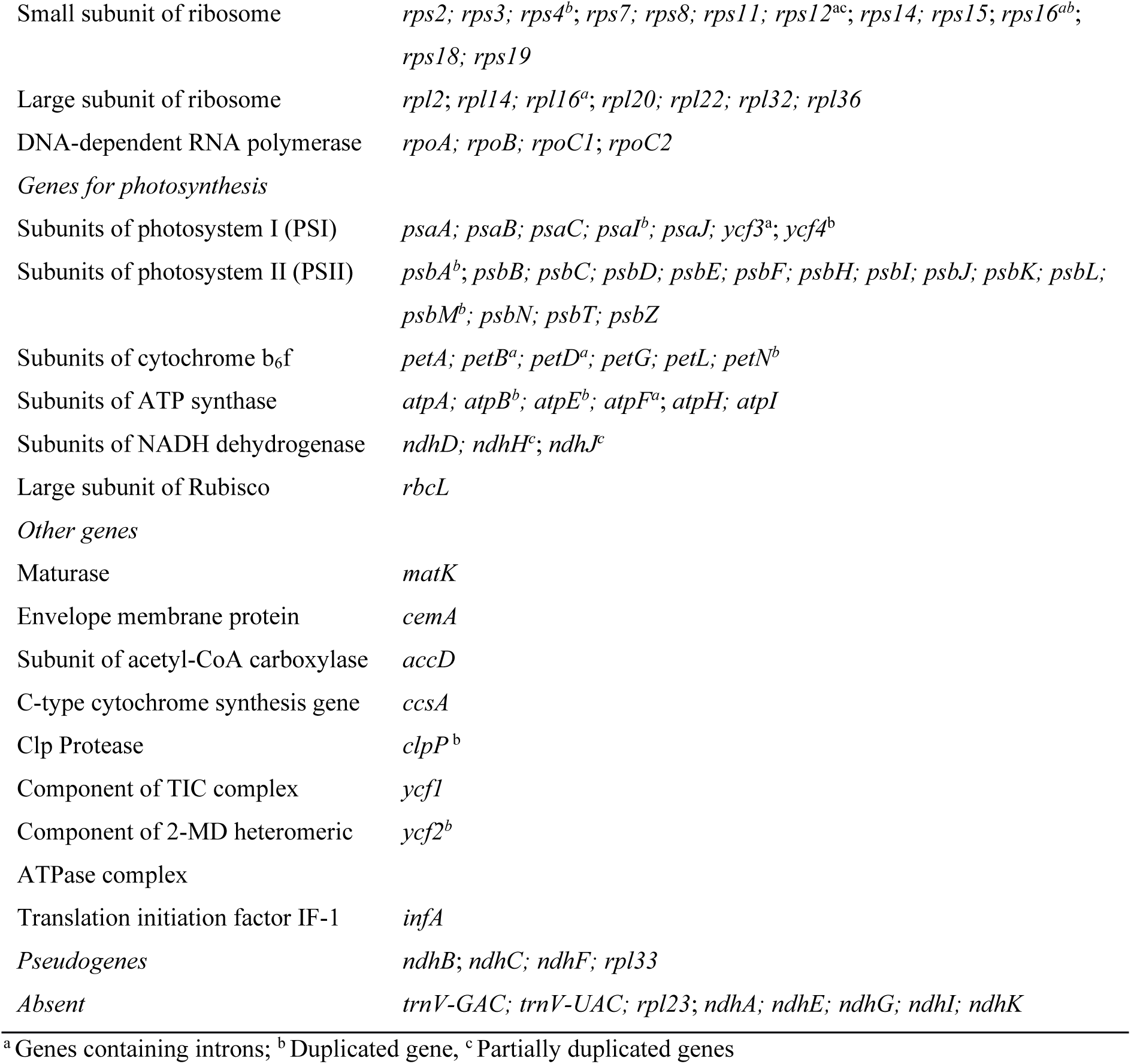
List of genes identified in the plastome of *Cereus jamacaru* subsp. *jamacaru*.

**Table 2.**
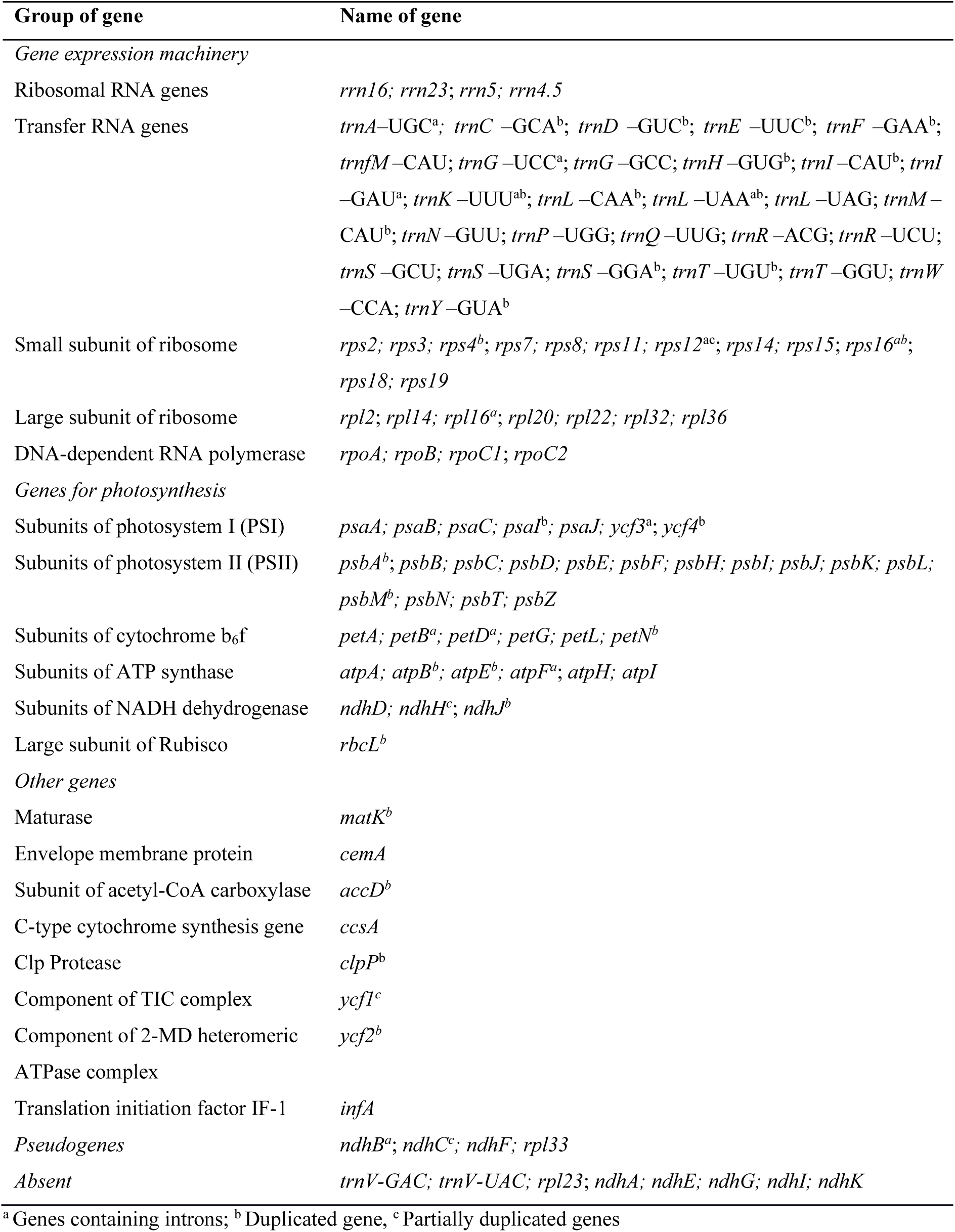
List of genes identified in the plastome of *Cereus hildmannianus* subsp. *hildmannianus*.

**Table 3.** General features of plastomes within Cactaceae.

| Species | Subfamily | Size<br>(bp) | LSC<br>(bp) | SSC<br>(bp) | IR<br>(bp) | Number<br>of genes | GC<br>(%) | GenBank |
| --- | --- | --- | --- | --- | --- | --- | --- | --- |
| *Cereus*<br>*hildmannianus* | Cactoideae | 141,600 | 53,310 | 24,320 | 31,955 | 101 | 36.6 | PQ997910 |
| *C. jamacaru* | Cactoideae | 141,884 | 53,632 | 23,778 | 32,237 | 101 | 36.6 | PV032414 |
| *C.*<br>*fernambucensis* | Cactoideae | 104,273 | N/A | N/A | N/A | N/A | N/A | OL397055.1 |
| *Melocactus*<br>*glaucescens* | Cactoideae | 129,806 | 51,465 | 19,097 | 29,622 | 98 | 36.5 | OK298499 |
| *M. ernestii* |  |  |  |  |  |  |  |  |
| *Rhipsalis teres* | Cactoideae | 122,389 | 81,397 | 24,016 | 8,488 | 99 | 36.7 | MT387452 |
| *Rhipsalis*<br>*baccifera* | Cactoideae | 122,333 | 81,459 | 23,531 | 8530 | 101 | 36.7 | MT821847 |
| *Selenicereus*<br>*undatus* | Cactoideae | 133,326 | 68,256 | 21,716 | 21,677 | 99 | 36.4 | NC_053698 |
| *Carnegiea*<br>*gigantea* | Cactoideae | 113,064 | - | - | - | 99 | 36.7 | NC_027618 |
| *Lophocereus*<br>*schottii* | Cactoideae | 113,204 | - | - | - | 99 | 36.5 | NC_041727 |
| *Mammillaria*<br>*albiflora* | Cactoideae | 110,789 | 78,380 | 31,061 | 674 | 96 | 36.4 | MN517610 |
| *M. pectinifera* | Cactoideae | 108,561 | 72,273 | 29,744 | 772 | 95 | 36.4 | MN519716 |
| *M. crucigera* | Cactoideae | 115,505 | 71,565 | 29,418 | 7,261 | 96 | 36.3 | MN517613 |
| *M.*<br>*huitzilopochtli* | Cactoideae | 115,886 | 71,997 | 29,401 | 7,244 | 97 | 36.3 | MN517612 |
| *M. solisioides* | Cactoideae | 115,356 | 71,690 | 29,238 | 7,214 | 95 | 36.4 | MN518341 |
| *M. supertexta* | Cactoideae | 116,175 | 72,240 | 29,445 | 7,245 | 97 | 36.4 | MN508963 |
| *M.*<br>*zephyranthoides* | Cactoideae | 107,343 | 71,811 | 7,281 | 14,126 | 95 | 38.5 | MN517611 |
| *Opuntia*<br>*quimilo* | Opuntioideae | 150,347 | 101,475 | 4,115 | 22,392 | 109 | 36.6 | MN114084 |

Of the 101 unique genes identified, 69 of them being protein-coding, 28 tRNAs, and 4 rRNAs (**Tables 1, 2**). Introns were identified in 11 genes, of which 6 are protein-coding genes and 5 are tRNAs. Two introns were identified in the *ycf3* gene, and the loss of one intron was observed in *clpP*, *rpl2*, and *rpoC1*. Thirteen tRNA and 11 protein-coding genes were duplicated. Partial duplications were observed for *rps12* and two *ndh* genes (*ndhH* and *ndhJ*; **Tables 1, 2**).

The *Cereus* plastome suffered a massive loss of 8 genes (*trnV-GAC, trnV-UAC, rpl23, ndhA, ndhE, ndhG, ndhI,* and *ndhK*), along with putative pseudogenization of 4 genes (*ndhB, ndhC, ndhF,* and *rpl33*). The determination of the *ndh* genes and *rpl33* gene as putative pseudogenes was based on the lack of initiation and/or termination codons, as well as premature stop codon resulting in non-functional fragments of the genes in question. Therefore, these genes can be identified the plastome, but its functionality is compromised.

The dot-plot analysis performed in MUMmer confirmed synteny between the plastome sequences and respective genomic regions (**Supplementary Figure S1**), and this result was also corroborated by whole genome alignment performed in MAUVE (**Supplementary Figure S2**). After confirming the synteny between *Cereus hildmannianus* subsp*. hildmannianus, C. jamacaru,* and *M. glaucescens* in NUCmer and MAUVE, we extended the analysis performed in Dalla-Costa et al. 2021 with our plastome data, exploring in detail the structural features evinced by the linear plastome maps drawn in OGDRAW webserver [18] (**Figure 4**). When compared to the canonical angiosperm plastome (*Portulaca oleracea*), the plastomes of *Cereus* had a large segment of the LSC presented a different arrangement. A primary translocation and inversion were identified for the region comprised between *trnQ*-*UUG* and *ycf4* genes (B, C, D, E, H, I, and J; **Figure 4**). Then, parts of this large segment had its direction reset after a secondary inversion (C, D, H, and I; **Figure 4**). Regarding gene losses in the *Cereus* plastomes, the genes *ndhK, trnV*-*UAC*, and *rpl23* were lost from the LSC (**losses 1, 2 and 3; Figure 4**); *trnV*-*GAC* was lost from the IRs (**loss 4; Figure 4**); and *ndhE*, *ndhG*, *ndhI*, and *ndhA* (**loss 5; Figure 4**), usually present in angiosperms presenting non-specialized lifestyles, were lost from the SSC. The genes *ndhF* (**loss 6; Figure 4**) and *trnA*-*UGC* (**loss 7; Figure 4**), absent in *M. glaucescens* plastome, are conserved in both *Cereus* species analyzed here (**Figure 4**). Concerning the structural rearrangement in the SSC, the rRNA genes were translocated from the IRs in *P. oleracea* to the SSC region (**Figure 4**).

**Figure 4.**
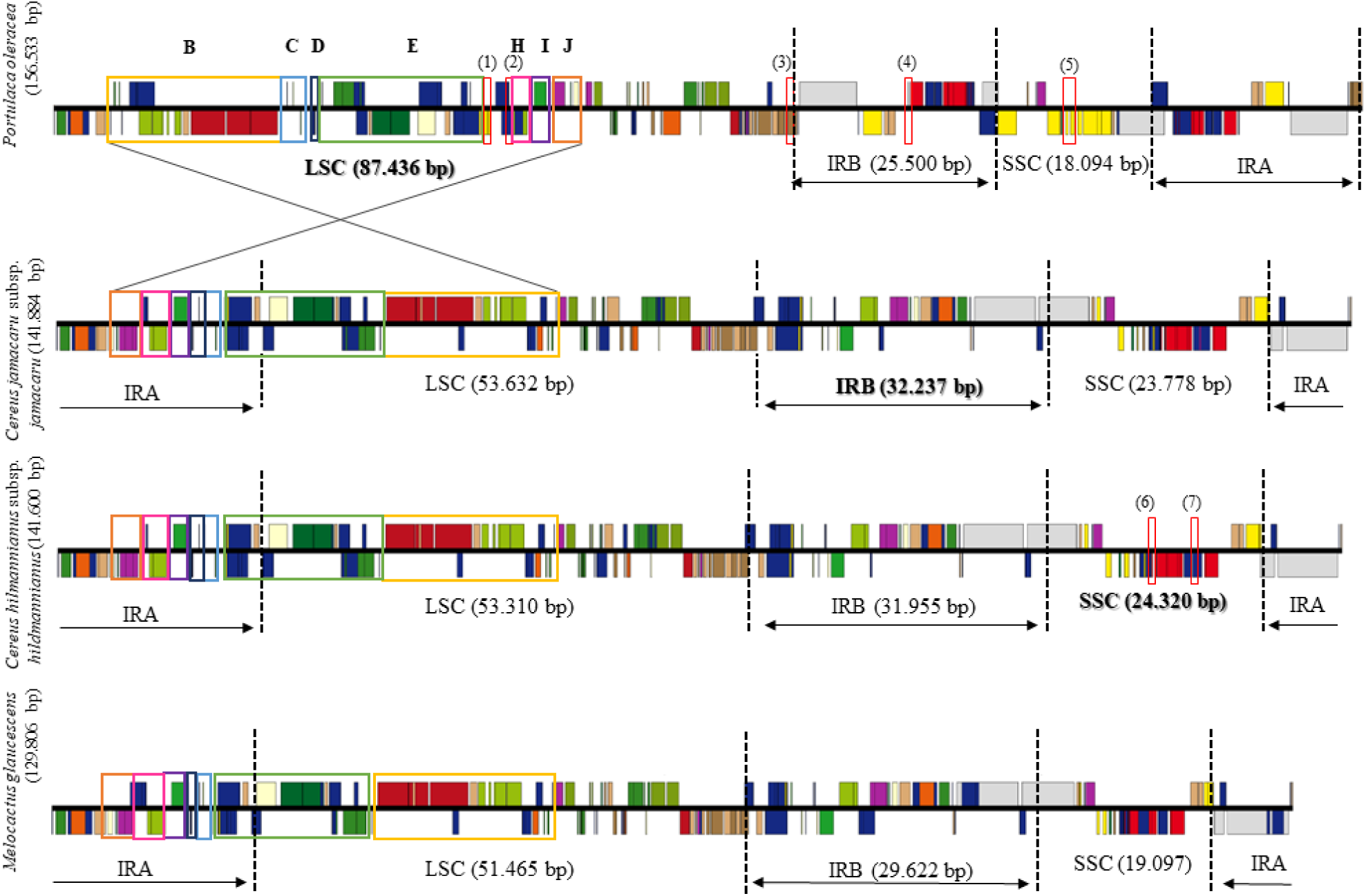
Gene content and order comparison between *Cereus hildmannianus subsp. hildmannianus, C. jamacaru* subsp. *jamacaru, Melocactus glaucescens* (subfamily Cactoideae) and *Portulaca oleracea* as a reference species for the typical angiosperm plastome. After confirming synteny between the *Cereus* species and *M. glaucescens* in NUCmer and MAUVE, we extended the analysis performed in Dalla-Costa et al. 2021 with our plastome data, exploring in detail the structural features evinced by the linear plastome maps drawn in OGDRAW webserver (Greiner et al. 2019). The colored blocks represent conserved regions across the genomes. Solid lines conecting the colored blocks in *P. oleracea* to those of *C. hildmannianus* show inversion and translocation of the gene order. The rearrangement events are described as follows: translocation and primary inversion of blocks **B**, **C**, **D**, **E**, **H**, **I**, and **J**; and secondary inversion of **B**, **C**, **D**, **H**, and **I**. Red squares indicate genes that were lost in the following plastome represented in line. (**1**) ndhK; (**2**) trnV-UAC; (**3**) rpl23; (**4**) trnV-GAC; (**5**) ndhE, ndhG, ndhI, and ndhA; (**6**) ndhF; and (**7**) trnA-UGC. The largest genomic regions between Cactoideae species are in **bold** characters, representing the expansion or retraction of the Large Single Copy (**LSC**), Small Single Copy (**SSC**), and Inverted Repeats A/B (**IRA/IRB**, which are identical in size and content).

### Codon Usage

The two tRNA genes: *trnV-GAC* and *trnV-UAC*, which encodes for the valine amino acid, are absent from both *Cereus*. We analyzed the codon usage frequency of these genes in *Cereus jamacaru* and *Cereus hildmannianus* and compared within the tribe Cereeae (**Figure 5**) and farther in the subfamily level, comparing the average of each tribe within Cactoideae (**Supplementary Figure S3**). The amino acid valine presented the frequencies of 0.14 for *C. jamacaru* and 0.18 for *C. hildmannianus* for the GTG codon (average for the subfamily: 0.14); 0.38 and 0.29 for GTA codon (average for the subfamily: 0.37); 0.35 and 0.32 for GTT (average for the subfamily: 0.36), and 0.12 and 0.21 for GTC codon (average for the subfamily: 0.13). *Cereus hildmannianus* presented above-average fractions for GTC and GTG codons, and lower fraction for GTA codon compared to the other members of the tribe (red arrows; **Figure 5**). The codon usage bias comparison between the Cactoideae tribes showed that the difference previously cited for *C. hildmannianus* is attenuated and that the fractions of both *Cereus* altogether are like other Cactoideae representatives, even in the absence of the tRNA genes encoding valine (**Supplementary Figure S3**).

**Figure 5.**
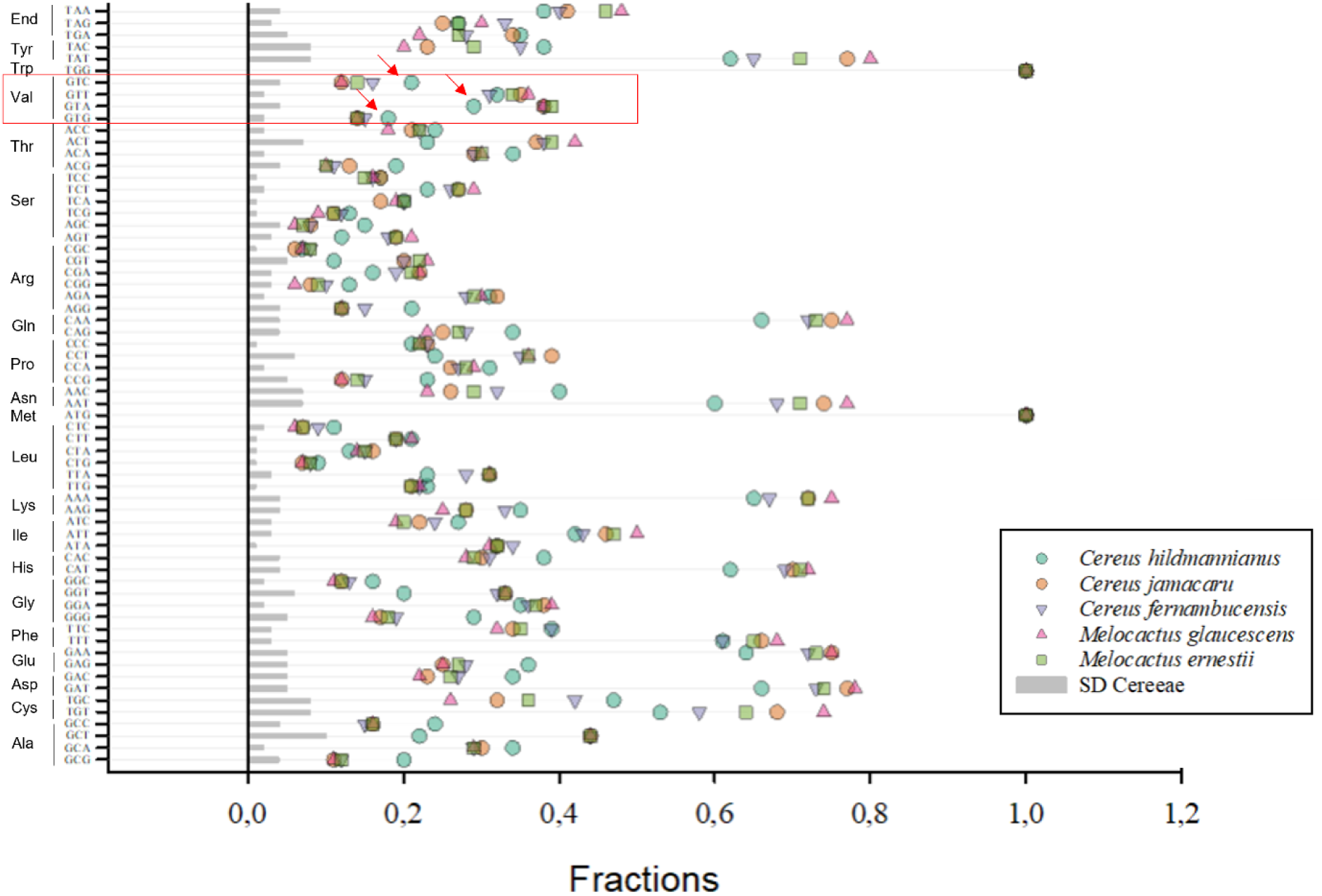
Frequency and standard deviation (SD) of the codon usage of species belonging to the Cereeae tribe: *Cereus hildmannianus subsp. hildmannianus, C. jamacaru* subsp. *jamacaru, C. fernambucensis, Melocactus glaucescens,* and *M. ernestii* plastidial protein-coding genes. The highlighted section corresponds to the codons for the amino acid valine, that had its respective tRNA genes (*trnV-GAC* and *trnV-UAC*) lost from *Cereus hildmannianus subsp. hildmannianus* and *C. jamacaru* subsp. *jamacaru* plastomes. The red arrows indicate outstandingly higher or lower values for *Cereus hildmannianus* subsp*. hildmannianus*.

### RNA editing sites in protein-coding genes of *Cereus* species and comparison to other cacti

We identified 8 editing sites in three genes that are exclusive of *Cereus*. *Cereus hildmannianus* harbors all of them (*rpoC1* – positions 391 and 540; *rpoC2* – 734 and 748; and *rps16* – 19, 40, 45, and 99), while *C. jamacaru* presents the first four. Of the 27 protein-coding genes analyzed here, we predicted 29 editing sites distributed in 15 genes for *C. hildmannianus,* and 24 editing sites distributed in 14 genes for *C. jamacaru* (**Supplementary Table S3**). Among them, the *rps16* gene presented editing sites (4) only for C. *hildmannianus*, and the *rpoC1* gene presented editing sites for and. The direct (C to U) editing sites were predicted to occur at the first (34.48% in and 37.5% in) and second (65,52% in and 62.5% in %) positions. Out of the 29 edition sites predicted for *C*. *hildmannianus,* 16 (55.17%) changed the polarity of the amino acid from polar to non-polar and four (13.79%) from non-polar to polar. Concerning the initially positive charged amino acid, one (3.45%) changed to non-polar (R to W); and four (13.79%) changed to polar. Four (13.79%) editions conserved the polarity of the amino acid (non-polar). Similarly, in *C. jamacaru,* of the 24 predicted editing sites, 8 (33.33%) changed the polarity of the amino acid from polar to non-polar, and two (8.33%) from non-polar to polar. Concerning the initially positive charged amino acid, all four (16.67%) changed to polar. Three (12.50%) editions conserved the polarity of the amino acid (non-polar).

Concerning the editing sites shared by both *Cereus* (24), 6 are shared with all Cactoideae species analyzed here (*atpA* (424), *petB* (140), *psbL* (1), *rpoB* (158 and 189), and *rps14* (50)). Two sites were shared by all Cactoideae except for *Mammillaria albifora*, who has a T fixed instead of edited (in genes/positions *atpA* (305) and *rps2* (83)). There was one editing site in *rpoB* gene shared by the subfamily, except for a few *Mammillaria* species (tribe Cacteae): position 184, which presented TTA (L) in *M. supertexta, M. pectinifera, M. crucigera, M. huitzilo-pochtli*, and *M. solisioides*. In the other species of the subfamily, the edition was predicted as changing from TCA (S) to TTA (L). *Mammillaria* also did not share the editing site occurring in *rpoA* gene (position 28) but did also present TAT (Y) which was, as in the previously described case, the final amino acid for the whole subfamily (that changed from CAT (H) to TAT (Y).

The editing sites predicted for *atpB* (pos. 140), *rpoC1* (14 and 210) were conserved in Cactoideae, except for *Melocactus glaucescens* (tribe Cereeae)*, Rhipsallis baccifera* (Rhipsalideae), and *Selenicereus undatus* (Hylocereeae). On the other hand, these three species were the only ones to share two editing sites with *Cereus hildmannianus* – *rpl20* (83), and *rpoC2* (498) – the latter being absent in *C. jamacaru.* The editing site at position 26 for the *psbF* gene (TCT (S) to TTT (F)) was shared between the two *Cereus* and *Carnegia gigantea* (Echinocereeae)*, Lophocereus schottii* (Echinocereeae)*, M. glaucescens, R. baccifera,* and *S. undatus*; the other species has a T fixed (TTT (F)). Three editing sites were conserved in Cereeae: *rpl20* (100), *rpoB* (158), and *rps8* (48).

### Single Sequence Repeats (SSRs) and Tandem Repeats mapping in the plastomes of *Cereus*

Of the 191 SSRs identified in *C. hildmannianus,* 137 were classified as mono-, 38 di-, 4 tri-, 7 tetra-, 3 penta-, and 1 hexapolymers, in addition to 1 compound SSR. The LSC comprised 86 (45.03%) SSRs; the IRB, 65 (34.03%); and the SSC, 40 (20.94%) (**Supplementary Table S4**). Regarding the region in the plastome, 97 (50.79%) SSRs were identified in intergenic spacers (IGS), 57 (29.84%) SSRs in coding sequences (CDS), and 37 (19.37%) in introns. Intergenic spacers that stand out regarding the higher SSR content (=3) are: *trnS-UGA/psbC, trnQ-UUG/cemA, petA/psbJ, rpl33/rps18, rps8/rpl14, rps4/trnT-UGU*, *trnC-GCA/trnE*-*UUC*, *accD*/*psaI*, *ccsA*/*ndhD*, *psaC*/*ndhH*-fragment, and *ndhH*-fragment/*rpl32*. Coding sequences where SSRs were more abundant were those in the genes *rpoB* (3), *rpoC2* (8), *petA* (2), *rps18* (2), *rpoA* (2), *accD* (2), *ycf4* (2), *ycf2* (6), *ycf1* (16), and *rpl32* (2); other genes presented only one SSR (**Supplementary Table S4**). The SSRs located in introns are spread across 10 genes, among protein-coding and tRNA genes (*ycf3*: 5; *atpF*: 7; *trnF-UCC:*2; *petB*: 2; *petD:* 4; *rpl16*: 2; *trnL-UAA:* 6; *rps16:* 3; *trnK-UUU:* 5; and *trnI-GAU*: 1).

Similarly, *C. jamacaru* contains 192 SSRs, being 139 mono-, 37 di-, 4 tri-, 7 tetra-, 3 penta-, and one hexapolymers, and one compound SSR (**Supplementary Table S5**). Eighty-eight (45.83%) SSRs were in the LSC region, 64 (33.33%) in the IRB, and 40 in the SSC (20.83%). Out of the 192 SSRs, 95 (49.48%) were found in intergenic spacers, 61 (31.77%) in coding regions, and 34 (17.71%) in introns. The SSRs found in intergenic spacers are mainly distributed in the regions: *ycf3/psaA, trnS-UGA/psbC, trnQ-UUG/cemA, petA/psbJ, rpl33/rps18, rps4/trnT-UGU, trnC-GCA/trnE-UUC, accD/psaI, ccsA/ndhD,* and *ndhH-fragment/rpl32,* presenting 3 SSR each. Additional information on IGS regions comprising less than 3 SSRs is available in **Supplementary Table S5**. Coding sequences where SSRs were more abundant were those in the genes *rps14* (1), *trnS-UGA* (1), *rpoB* (3), *rpoC1* (1), *rpoC2* (8), *rps2* (1), *trnG-UUC* (2), *clpP* (1), *cemA* (1), *petA* (2), *rps18* (3), *rpl20* (1), *rpoA* (2), *rpl22* (1), *rbcL* (1), *atpB* (1), *accD* (2), *ycf4* (2), *ycf2* (6), *ycf1* (16), *rps15* (1), *rpl32* (2), and *rrn23* (1).The SSRs located in introns are spread across 9 genes, as follows: *ycf3* (5), *atpF* (7), *petB* (2), *petD* (3), *rpl16* (2), *trnL-UAA* (6), *rps16* (3), *trnK-UUU* (5), and *trnI-GAU* (1).

We identified 50 tandem repeats (TRs) for the plastome of *C. jamacaru* and 44 for *C. hildmannianus hildmannianus* (**Supplementary Tables S6 and S7**, respectively). The first presented 24 (48%) tandem repeats in protein-coding regions, 19 (38%) in intergenic spacers, (6%) three in introns, two (4%) in rRNA genes, and two (4%) that were partly in intergenic spacer region, and partly in protein-coding region. Regarding the regions in the plastome that presented higher nucleotide repeat regions, 26 (52%) were found in the LSC, 14 (28%) in the SSC, and 10 (20%) in the IRB (the IRA was excluded to avoid data redundancy). Seven (14%) TRs were found in the *ycf1* gene and two (4%) in the *ycf2*.

Likewise, the 44 tandem repeats identified for *C. hildmannianus* were distributed as follows: 14 (31.82%) in protein-coding regions, 16 (36.36%) in intergenic spacers, six (13.64%) in introns, two (4.55%) in rRNAs, and five (11.36%) comprised in between a CDS and intergenic spacer. Out of the 44, twenty (45.45%) were in the LSC, 13 (29.55%) in the SSC, and 11 (25%) in the IRB.

The ribosomal S18 gene bears 9 tandem repeats in *C. jamacaru,* a strinkingly higher number when compared to TRs observed in related genes (S16 and S19 presented two TRs each; and S3, only one). Differently, the genome of *C. hildmannianus* presented only two TRs for the *rps18* gene. Since this is not a highly divergent gene, i.e., presents usually a conserved structure and content across Cactaceae, we attribute this result to an insertion found in this gene. It is larger in *C. jamacaru* than in *C. hildmannianus* and was not described for other cacti to date. The complete list of SSR loci and tandem repeats is available in the **Supplementary Tables S4, S5, S6, and S7**).

### Unusual insertion in the protein-coding gene *rps18*

We identified, for both species of *Cereus*, an extension in the protein-coding gene *rps18*. This gene was 1.308 and 927 bp long for *C. jamacaru* and *C. hildmannianus* respectively. In *C. jamacaru* the insertion was 851 bp long (**Figure 6**), whereas in *C. hildmannianus* it was 444 bp long. Although the sequence of is way larger than the others, the insertion itself is partly shared between both *Cereus* (**Figure 6**). Interestingly, the closest species (i.e. belonging to Cereeae) do not share of this insertion; instead, a species of Trichocereeae tribe (namely *Gymnocalycium saglionis*) and one of Notocacteae (*Frailea castanea*) presented higher similarity to *Cereus rps18* sequence than *Melocactus glaucescens* or *M. ernestii,* both belonging to Cereeae and sharing evolutionary features.

**Figure 6.**
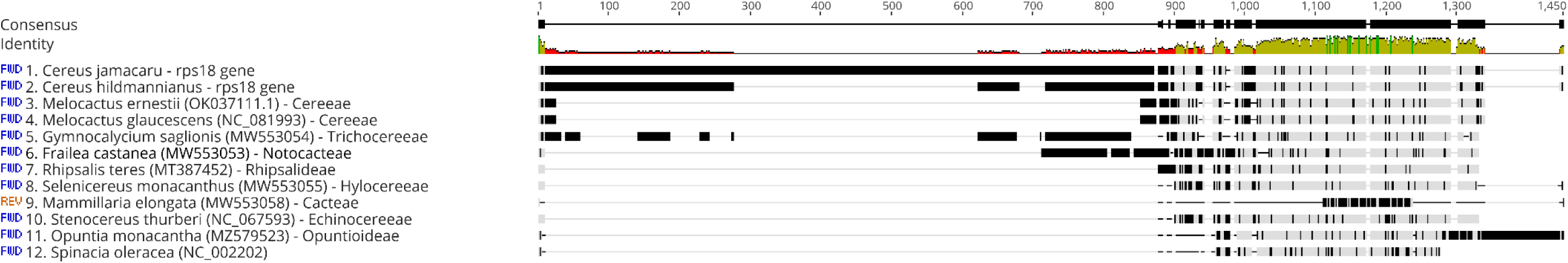
MAFFT alignment of the *rps18* gene in *Cereus hildmannianus* subsp.*hildmannianus, C. jamacaru* subsp. *jamacaru,* and species belonging to Cactoideae tribes and *Spinacia oleracea*. The tribes are indicated in each sequence, after the accession number.

The first round of SSR analysis performed in MISA did not identify repeated regions within this gene. When performing this analysis under default parameters (i.e., more relaxed than the ones set initially) of the webserver, it predicted a compound repeated region (in *C. jamacaru*, 34 bp long: (T)10aataaaattt(TA)7; in *C. hildmannianus*, 43 bp long: (T)11aataaaattt(TA)11;) in the adjacent intergenic region comprised between the *rpl33* and the *rps18* genes for both *Cereus* (SSR number 14; **Supplementary Tables S4 and S5**). In addition to the repeat region identified previously in MISA for the intergenic region between the S18 and L33 genes, the tandem repeat analysis performed in Phobos v. 1.0.6 identified 8 repeat regions of varied size and content within the protein-coding sequence of the *rps18* gene (**Supplementary Tables S6 and S7**).

### Phylogenetic reconstruction

The phylogenetic tree reconstructed by Maximum Likelihood shows a log-likelihood of -104642.5982, AICc of 209613.00, and represents 58 *taxa* with six partitions, 29824 total sites and 0% missing data after careful visual curation and running the gap removal script developed here for improved refinement of the gene alignments (**Figure 5, Supplementary Figure S4, Supplementary Script S1**).

The relationships within Cactaceae are strongly supported with many nodes at 100% bootstrap for the clades Cereeae, Rhipsalideae, Hylocereeae, Cacteae, Opuntioideae, as well as between Cactaceae and Portulacaceae and outgroups used here (**Supplementary Figure S4 and Supplementary Table S2**). Only a few clades present lower support due to non-zero internal branches, possibly indicating polytomies according to the logfile (46% in *Ariocarpus*; 46% in *Selenicereus*; 44% in the basal split with *Halophytum ameguinoi* and *Anredera cordifolia*; **Supplementary Figure S4**).

The genus *Cereus* is supported as a monophyletic group within its tribe with a bootstrap = 100, as well as the other relationship in Cereeae, with exception of *Gymnocalycium saglionis* (BS=65). The sister tribes Rhipsalideae and Hylocereeae are monophyletic and well supported (BS=100) in our reconstruction, despite the low support of *Selenicereus undatus* (Hylocereeae). The weak nodes need additional data for deeper investigation.

Cacteae, the most species-rich tribe of Cactaceae, is well supported and monophyletic, with its relationships largely presenting a high bootstrap value. The bootstraps equal 100 in all the genus nodes. The species *Mammillaria zephyranthoides* and *M. supertexta* presented bootstraps of 94 and 72, respectively. *Ariocarpus*, although well-supported as a genus, presented low support values within its species (**Figure 7**, **Supplementary Figure S4)**. The sister clade comprising *Ferocactus, Leuchtenbergia,* and *Echinocactus* is well-supported and presents little variation of BS support among its genera. Opuntioideae stands out as the clade that first diverged in Cactaceae, being well-supported and monophyletic as well.

**Figure 7.**
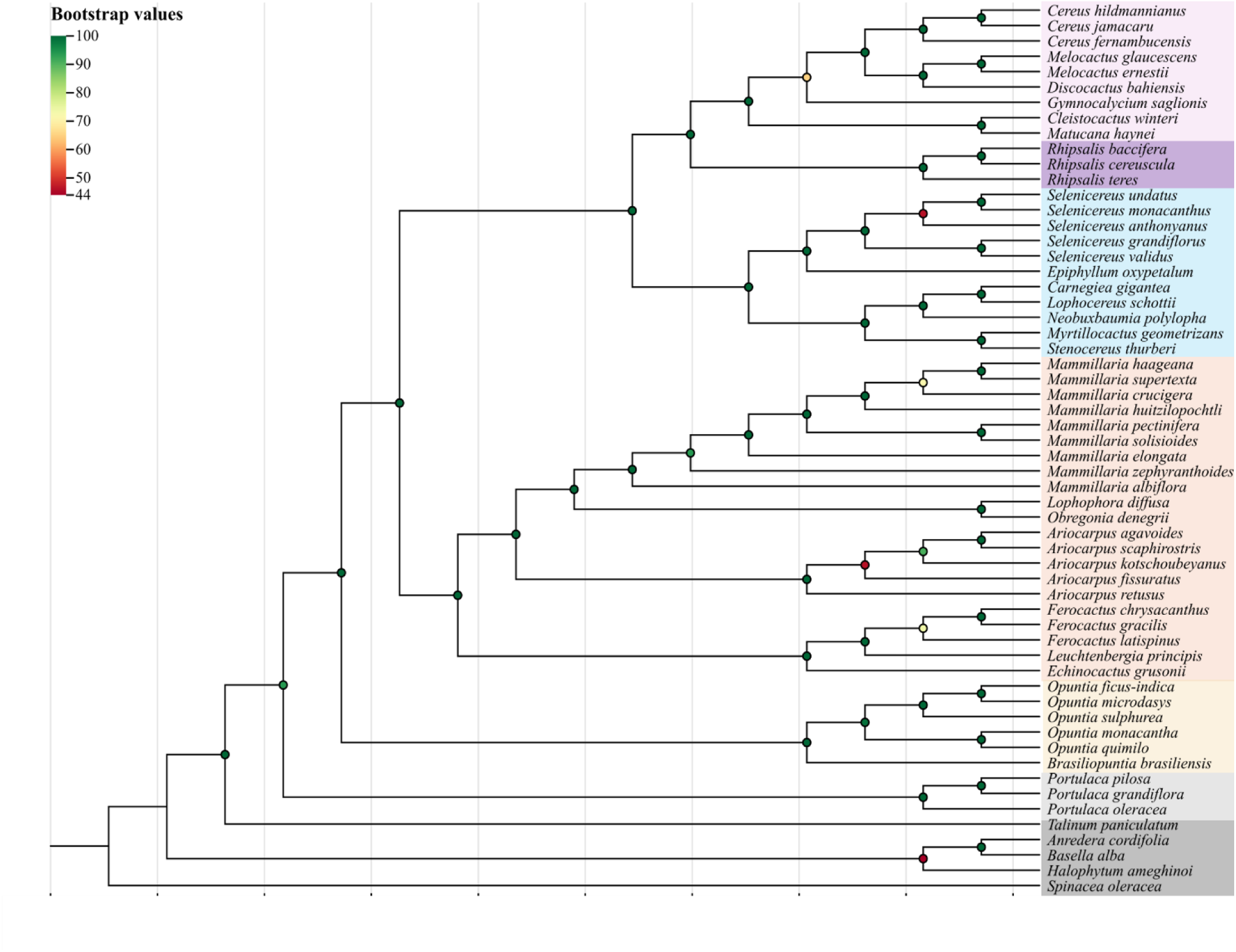
Phylogenetic tree reconstructed by maximum-likelihood based on 45 shared protein-coding genes of 58 cacti species. The bootstrap values are color-coded, green circles above 80, yellow between 80 and 60, and red below 60, as indicated in the legend in the top-left. Species are grouped by tribe, indicated in the right. Colored blocks refers to each tribe.

Among the outgroups, *Talinum paniculatum* (Talinaceae; **Supplementary Table S2**) was closer to Cactaceae and Portulacaceae, presenting bootstrap of 94. Basellaceae, represented by *Anredera cordifolia* and *Basella alba,* and *Hallophytaceae,* represented by *Halophytum ameghinoi*, formed a sister group in which the latter family diverged earlier. *Spinacea oleracea* was used as canonical sequence for rooting the tree.

## Discussion

### General features of *Cereus* plastomes

Cactaceae stands out as a highly speciose family, with a myriad of lifeforms, and spanning a broad range of occurrence throughout the Neotropics, except for *Rhipsalis baccifera* that occurs in the old world [29]. As such, not only the family, but also Cactoideae holds an intriguing evolutionary history in its plastid genomes, presenting important events such as subfamily-specific rearrangements, IR expansions and retractions, gene losses and pseudogenization [30–36].

As demonstrated recently [31, 32], Cereeae representatives present usually between 120-130 kbp, unlike other basal Cactaceae subfamilies (*e.g.* Opuntioideae – with *Opuntia quimilo* presenting a plastome as large as 150.347 bp). The plastomes of *Cereus,* however, are among the largest observed for Cereeae to date (*C. jamacaru –* 141.884 bp; *C. hildmannianus* – 141.600 bp), being structurally very similar to the plastome of *Acanthocereus tetragonus,* which is larger in size (142.663 bp) [31].

Although there is clear evidence of variation of size within Cereeae, the structure of the plastid genomes of the species therein is usually similar, and such pattern was recently described as a determined type of plastome when compared to other tribes (“Type E”; [31]). The plastomes of the two *Cereus* analyzed here are highly syntenic, containing same gene content and order. Eight genes (*trnV-GAC, trnV-UAC, rpl23, ndhA, ndhE, ndhG, ndhI,* and *ndhK*) are absent from *Cereus* plastome, and other 4 genes (*ndhB, ndhC, ndhF,* and *rpl33*) are presumably pseudogenes (*i.e.*, have their structural signature in the plastome, but lack functionality).

The loss of the *rpl23* gene (which encodes for the L23 protein of the ribosomal 50S) is expected, since this gene tends to be degenerated or absent in multiple lineages within Cactaceae (*M. glaucescens*, *S. truncata, L. cruciforme, R. baccifera,* and *R. teres*) [29, 32–34] and other angiosperms [37–40]. The degradation of this gene may indicate that a substitute may be imported from the cytosol [38]. The loss of two tRNA genes (*trnV*-*UAC^Val^*and *trnV-GAC^Val^*) is discussed in the upcoming sections.

Loss and pseudogenization of *ndh* genes are often described for many angiosperms lineages, and this frequently degenerated set of NAD(P)H dehydrogenase genes can affect the management of photo-oxidative stress response, therefore compromising photosynthetic efficiency in terrestrial plants [41, 42]. Although the most accentuated losses of the *ndh* complex can be observed in heterotrophic lineages, where a decrease in photosynthetic efficiency won’t likely affect the organism’s survival, the extreme reduction of the *ndh* set is also related to species presenting CAM metabolism. The CAM metabolism compensates the *ndh* genes’ functionality by enhancing CO_2_ concentrations, limiting photorespiration, and therefore improving CO_2_ uptake by the organism [32, 33, 43, 44].

When considering organism development and survival, the presence or absence of *rpl33* in plant species seems to be arbitrary. This gene encodes the L33 protein of the ribosomal 50S subunit, playing an important role in protein synthesis during the early stages of plant development [45]. While its presence in some cacti species is likely explained by their slow-growing lifestyle (*M. glaucescens*)[32], the retention of this gene in fast-growing species is contradictory to the lifestyle explanation (*Hatiora salicornioides*, Badia et al., *in prep; R. baccifera, Schlumbergera truncata*) [29, 33]. Its absence, for its turn, is tolerated by some species (*R. teres*; *Lepismium cruciforme*) [33, 34], but known to be required under cold-stress conditions [45].

We identified expansion of the IRs at the expense of reduction of the large single copy, when considering the typical angiosperm plastome. The LSC is 53.632 bp large in *C. jamacaru* and 53.310 in *C. hildmannianus* subsp. *hildmannianus,* whereas in *P. oleracea* the LSC is 87.436 bp long. In turn, the IR in *P. oleracea* is only 25.500 bp long, versus 32.237 bp and 31.955 bp long in *C. jamacaru* and *C. hildmannianus* respectively. Genes are being imported to the inverted repeat region and naturally being duplicated. This indicates that these genes are potentially fundamental for fitness of these species.

The overall high synteny between *Cereus* plastids shows that the divergent environmental pressures across its sparse range of occurrence have not led to strong structural changes in the plastome, as the genus has a relatively recent divergence (∼2.6 million years ago) [12].

### The superwobbling phenomenon explains the lower codon usage fractions in *Cereus*

Our results show that two tRNA genes encoding for valine were lost from *Cereus* plastomes: *trnV-GAC* and *trnV-UAC.* Reverse genomics points out that the *trnV-GAC* can be absent without compromising cell viability, whereas *trnV-UAC* is essential for the same purpose [46]. Interestingly, *M. glaucescens* share the loss of these two *tRNA*s [32], and the loss of *trnV-UAC,* regarded as essential, is also common to *Discocactus bahiensis* and *M. ernerstii* [47] and to Cactoideae in a broader sense [48].

The wobbling rules proposed by Crick state that a minimum of 32 tRNAs are required to decode all codons, considering some flexibility at the third codon position to optimize codon-anticodon pairing [49]. Following the wobbling rules, these *tRNA*s are responsible for decoding four valine codons (GUA, GUC, GUG, and GUU). Under specific conditions, the wobbling rules are ruled out, giving place to an ongoing super wobbling mechanism. This mechanism enables a single tRNA to recognize all four codons in a degenerate codon box upon presence of an unmodified uridine at the wobble position, hence compensating an eventual reduction in the tRNA dataset [46, 50].

While *C. jamacaru* fractions for the four valine codons fit within the average of the tribe, *C. hildmannianus* presented above-average fractions for GTC and GTG codons, and lower fraction for GTA codon compared to the tribe (red arrows; **Figure 5**). The valine codon fractions observed for *C. hildmannianus* may indicate that a superwobbling is taking place in this species, especially if considered that this species inhabits a very distinct environment than typical Cereeae. Alternatively, higher fractions despite tRNA^val^ absence indicate that the product is being imported from the cytosol, although evidence that this mechanism occurs in plastids is still incipient.

### Differently shared RNA-editing sites between *Cereus* may indicate mutations shaped by different habitats

The RNA-edition prediction is often used as a tool to predict which regions of the plastid genome are susceptible to changing their basis to regulate gene expression under distinct conditions, including physiological stress [51].

The plastid genome of *C. hildmannianus* showed a higher frequency of putative RNA editions (29 in 15 genes) in comparison to *C. jamacaru* (24 in 14 genes). Considering all editions for both species, eight are exclusive of these species*. Cereus hildmannianus* harbors all of them, distributed in the genes *rpoC1*, *rpoC2,* and *rps16*, whereas its relative *C. jamacaru* shares four predictions, only in the *rpo* genes.

The higher frequence of the RNA editing predictions in *C. hildmannianus* are possibly related to the environmental constraints in the Pampa biome, where the thermal amplitude is greater than in Caatinga biome [52]. Considering that most of the Cactaceae occurrence is related to arid environments with high temperatures and low humidity, it is reasonable to assume that a cactus species under cold-stress regime may present adaptative mechanism in response to it. This has been observed in other plant groups under oxidative or temperature stress, where RNA editing seemed to contribute to the stabilization of the gene function [53, 54].

The *rps16* gene is known for being highly divergent, making it a valuable marker phylogenetic inferences along with *trnL-trnF* and *trnK-matK* regions [55]. Despite this variability, this gene holds essential functions even in heterotrophic *taxa,* indicating strong evolutionary signatures on the preservation of this gene over plant evolution [56]. The fast-evolving nature, associated with the fact that this gene is essential, facilitates the presence of RNA editing, a mechanism that may compensate for deleterious mutations and maintain functionality. The *rps16* gene belongs to the ribosomal protein gene category and plays a role in plastid ribosome assembly, a core function for gene expression for the plastid [57]. Therefore, we can infer that both the frequency and spatial distribution of RNA editing sites in *Cereus* likely reflect adaptive responses to ecological constraints.

### The high number of SSRs represents an important source of molecular markers aiming at conservation strategies

*Cereus hildmannianus* and *C. jamacaru* presented 191 and 192 SSRs, respectively (**Supplementary Tables S4 and S5**). The prevalence of SSRs is in the LSC, followed by the IR and SSC regions. The intergenic spacers presented a higher number of SSRs, followed by coding-regions and introns. The Tandem Repeats were also found in higher frequency in the LSC, but differently from the SSRs, the SSC presented higher frequency of TRs than the IR. A higher number of TRs was identified for the protein-coding regions, followed by the intergenic spacers and introns.

The regions of the plastome may differ as for the frequency of polymorphic regions. In species where the IR is not under deep changes in size and content, it was described as a higher frequency of polymorphic sites in the SSC region when compared to the other regions of the plastome [58]. This difference was, in the mentioned case, explained by the repair mechanism that would be sharper in the palindromic repeat regions than in single copy regions. Differently, in the plastomes investigated here we observed that the IRs presented the highest polymorphic frequency. This would therefore indicate a more relaxed repair mechanism in the IR regions, since the inverted repeats in Cactaceae tend to suffer great expansion, or retraction, or even complete loss [32, 43].

The intergenic regions are usually expected to harbor most of the polymorphic regions, but here, only the SSRs were described as higher frequency. Due to their overall higher mutation rates and faster evolution than coding-regions, which tend to be conserved, the intergenic regions are the ideal portions of the genome to study genetic divergence.

The intergenic regions are suitable regions to access when aiming at unveiling the genetic structure of a given species; and the genetic structure can therefore be used for a deeper understanding on the phylogeographic history of that species, phylogenetic relationships between species, or even conservation strategies. The intronic regions possess similar characteristics, but in our results, presented the lower proportion of SSRs when compared to other studies [32, 58]. Additionally, the high frequency of SSRs in coding regions observed for both *Cereus* is unusual, and that could be related to gene duplication in these species and to a relaxed repair mechanism. Likewise, the Tandem Repeats identified for both species

### Insertion in the *rps18* gene provides insights on the plastid DNA evolution in Cactaceae

The intron gain in plants is quite rare, but recent studies have been shedding light on the understanding of these regions (e.g. *rps18* presenting two introns in *Lonicera japonica,* [59]). Here, we detected an insertion on the ribosomal protein S18 gene in plastomes of *Cereus*. This gene is regarded as essential for plant development [60]; therefore, changes in its structure should not alter its functionality. In both *Cereus* the gene is functional, but divergent from other cacti sequences. When comparing to other Cactoideae species, the insertion is partially shared with species that theoretically do not present intron in this gene (**Figure 6**). Given its high frequency in plants, especially in plants presenting Crassulacean Acid Metabolism (e.g. orchids, [61]), it is reasonable to assume intron retention as a possible explanation for this event [62, 63].

Our genomic data provides useful insights about a potential alternative splicing event that may be occurring in the plastomes of the *Cereus* species studied here. Predictions based on structural genomic data presented here, such as intron presence prediction and single gene phylogeny, would be useful to access the evolution of this gene throughout the lineages. Additionally, the genomic data can guide further experimental validation and deeper investigation using transcriptome sequencing. Accessing the transcripts of this gene would cover in deeper level the complexity and tissue-specific nature of the ongoing splicing events. Functional genetics would offer important insights on which genes are being expressed relative to the distinct habitats in which they prevail.

### Phylogenetic reconstruction

*Cereus* is strongly supported within Cereeae, presenting bootstrap of 100% in all internal nodes, thus reinforcing the monophyly of *Cereus* and its close relationship with *Melocactus* and *Discocactus*. The closest relationship between *C. jamacaru* and *C. hilmannianus* likely suggests recent speciation or ongoing gene flow, a pattern observed in previous studies on Pleistocene diversification of the genus [64]. The basal branch of *C. fernambucensis* within Cereeae, however, may indicate a more ancient divergence of this species, which may be related to historical climate fluctuations [65].

Rhipsalideae is well supported, and Hylocereeae stands out by possessing one of the lowest bootstraps supports in our phylogenetic reconstruction (BS=46). The closest relationship between *Selenicereus undatus* and *S. monacanthus* likely indicates rapid radiation, a phenomenon previously reported for epiphytic cacti [44], that may be affecting the resolution of this section. The same applies to *Mammillaria haageana* and *M. supertexta* (tribe Cacteae; BS=72). In Cereeae, *Ariocarpus* displays as a monophyletic group, despite the low support (BS=46) observed. Other hypotheses known to affect Cactaceae that may be causing low support values are incomplete lineage sorting or hybridization [4]. Further studies addressing these species from other perspectives such as nuclear markers approaches would advance knowledge on this section, as is the case for recently published work on related species [47].

*Gymnocalycium saglionis* displays the longest branch length in Cereeae and moderate support (BS=65), suggesting that *G. saglionis* diverged earlier than *Melocactus*, *Discocactus* and *Cereus*, and has the most divergent sequence between the samples analyzed here. This result corroborates previous findings indicating that this genus often present high genetic divergence driven by adaptation to different ecological and environmental conditions [66]. Additionally, the clade Opuntioideae presented a highly supported monophyly, reinforcing its status a distinct lineage [1].

## Conclusions

Compared to the typical angiosperm plastome structure, both plastomes presented several distinct features within the Cactaceae, such as structural rearrangements, gene losses, and atypical gene duplications, which led these genomes to be amongst the largest of all Cactaceae described so far. We identified single sequence repeats (SSRs) and tandem repeats, which are fundamental for evaluating the genetic diversity of natural populations for conservation purposes, especially considering the historical exploitation of members of this family. The RNA editing sites predicted here provide insights on the eco-evolutionary aspects of the genus. We mapped a significant loss of most of the genes of the *ndh* complex, and the loss of three tRNA genes that, along with the codon usage results, indicate that such genes are being imported from the cytosol to the plastids or, alternatively, a superwobbling mechanism is likely in action. A possible intron retention may be in course for the gene *rps18*, given its difference from other related species. Our results bring a broader perspective not only on plastome evolution in the tribe Cereeae, but also on phylogenetic reconstructions for the suborder, that has long been discussed in the literature. Our findings emphasize the need for integrative methodologies that allow a broader comprehension of the evolutionary history of Cactaceae, thus combining morphological, ecological, and genomic data.

## Author contribution statement

Material preparation, data collection and analysis were performed by Clara C.V. Badia and Maria C. Silva. The first draft of the manuscript was written by Clara C.V. Badia. All authors read and approve the final manuscript. Marcelo Rogalski, Jeferson Fregonezi, Clara C.V. Badia and Maria C. Silva conceived and designed the research. Valter A. de Baura, Eduardo Balsanelli, Emanuel Maltempi de Souza, Fábio de Oliveira Pedrosa, Jéferson Fregonezi and Marcelo Rogalski contributed with reagents and materials.

## Acknowledgments

This research was supported by the National Council for Scientific and Technological Development, Brazil (CNPq - Grants 459698/2014-1, 310654/2018-1 and 436407/2018-3). Clara C.V. Badia acknowledges financial support from the Coordination for the Improvement of Higher Education Personnel (CAPES) through her doctoral studies, including CAPES-PrInt scholarship for studying abroad during part of her research. The authors also thank Cactário Guimarães Duque (CAGD – INSA/MCTI) for providing the plants used in this study, and Caio Crelier and Alenilson Rodrigues for kindly sharing the pictures of the species.

## SUPPLEMENTARY MATERIALS

**Supplementary Fig. S1.**
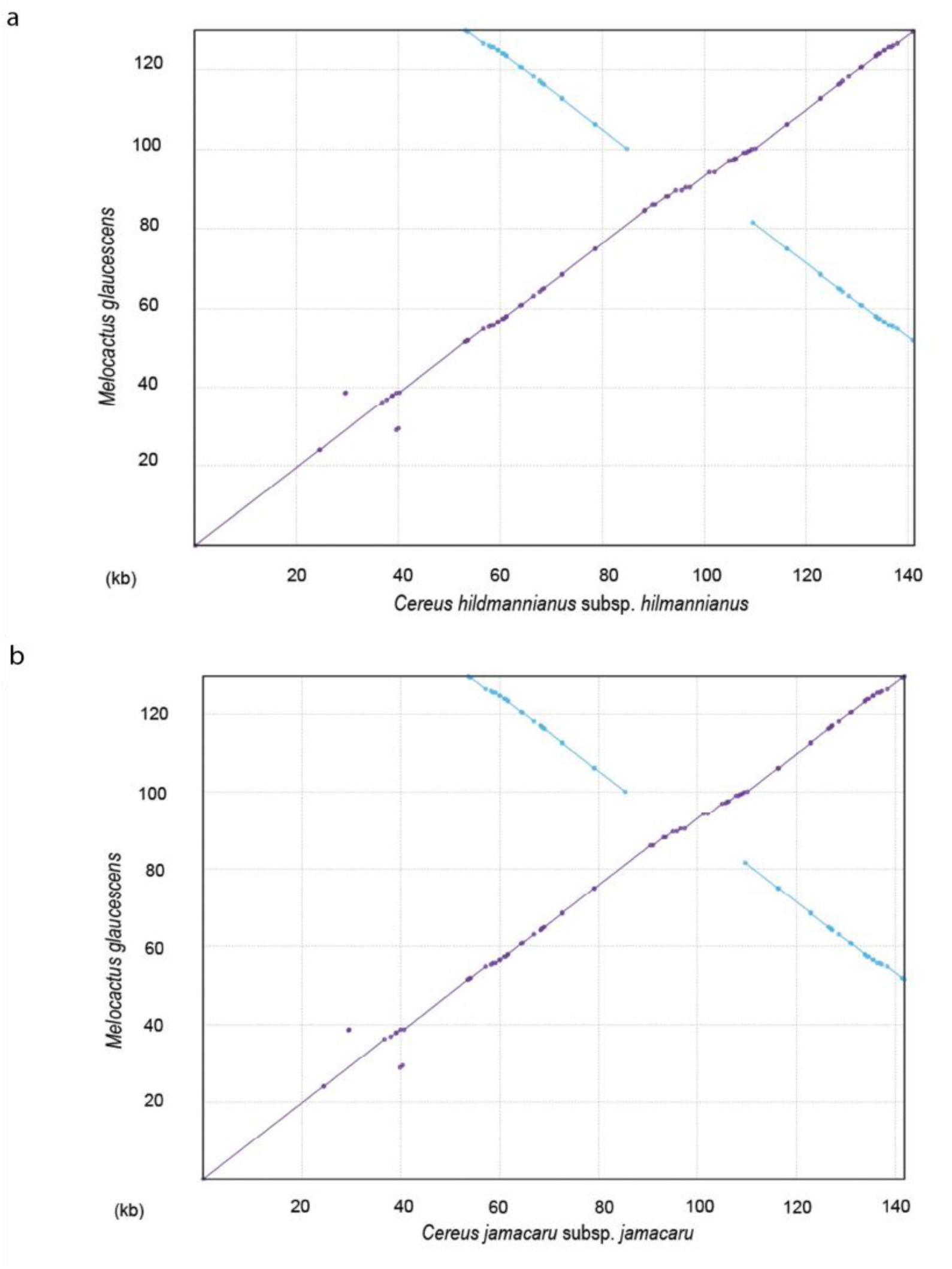
Dot-plot analysis comparing the complete plastomes of (**a**) *Cereus hildmannianus* subsp. *hildmannianus* and (**b**) *C. jamacaru* subsp. *jamacaru* to *Melocactus glaucescens*. The positive slope observed in the purple lines indicates sequences in the same direction, whereas the negative slope represented by blue lines indicate sequences with opposite direction.

**Supplementary Fig. S2.**
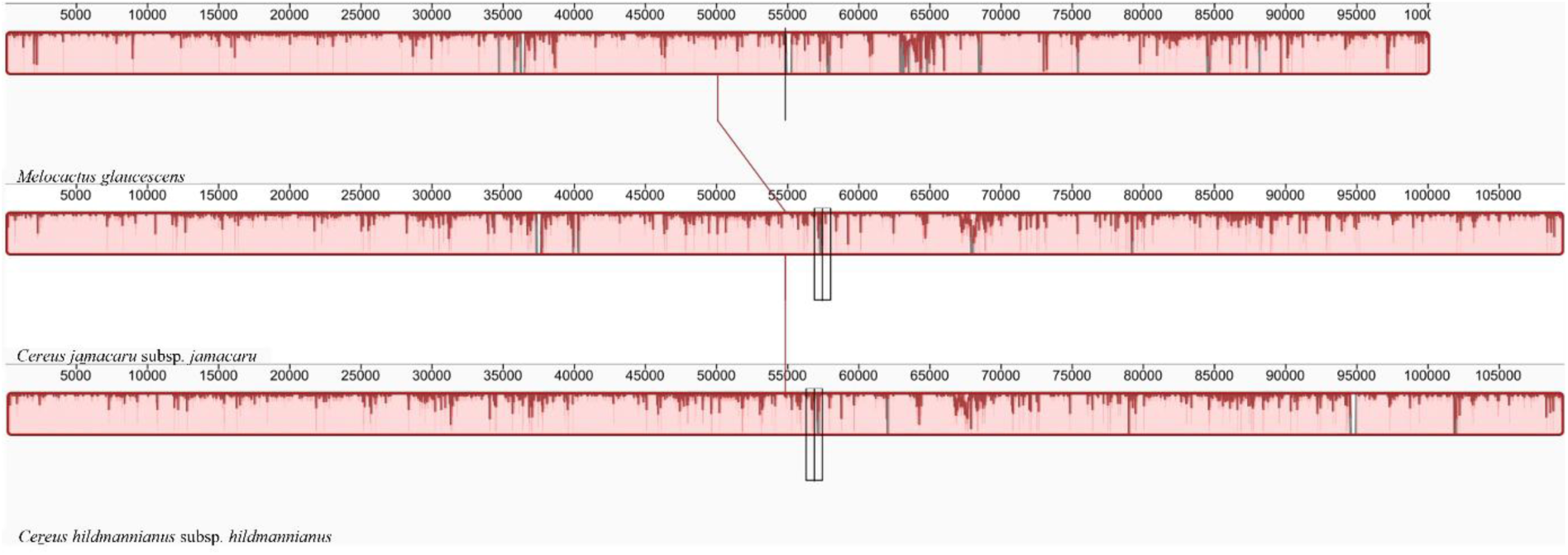
Multiple alignments comparing the plastomes of *Cereus jamacaru* subsp. *jamacaru* and *C. hildmannianus* subsp. *hildmannianus* to *M. glaucescens* (subfamily Cactoideae). The selected region of the plastomes includes both single copies and one inverted repeat (LSC, SSC, and IRB).

**Supplementary Fig. S3.**
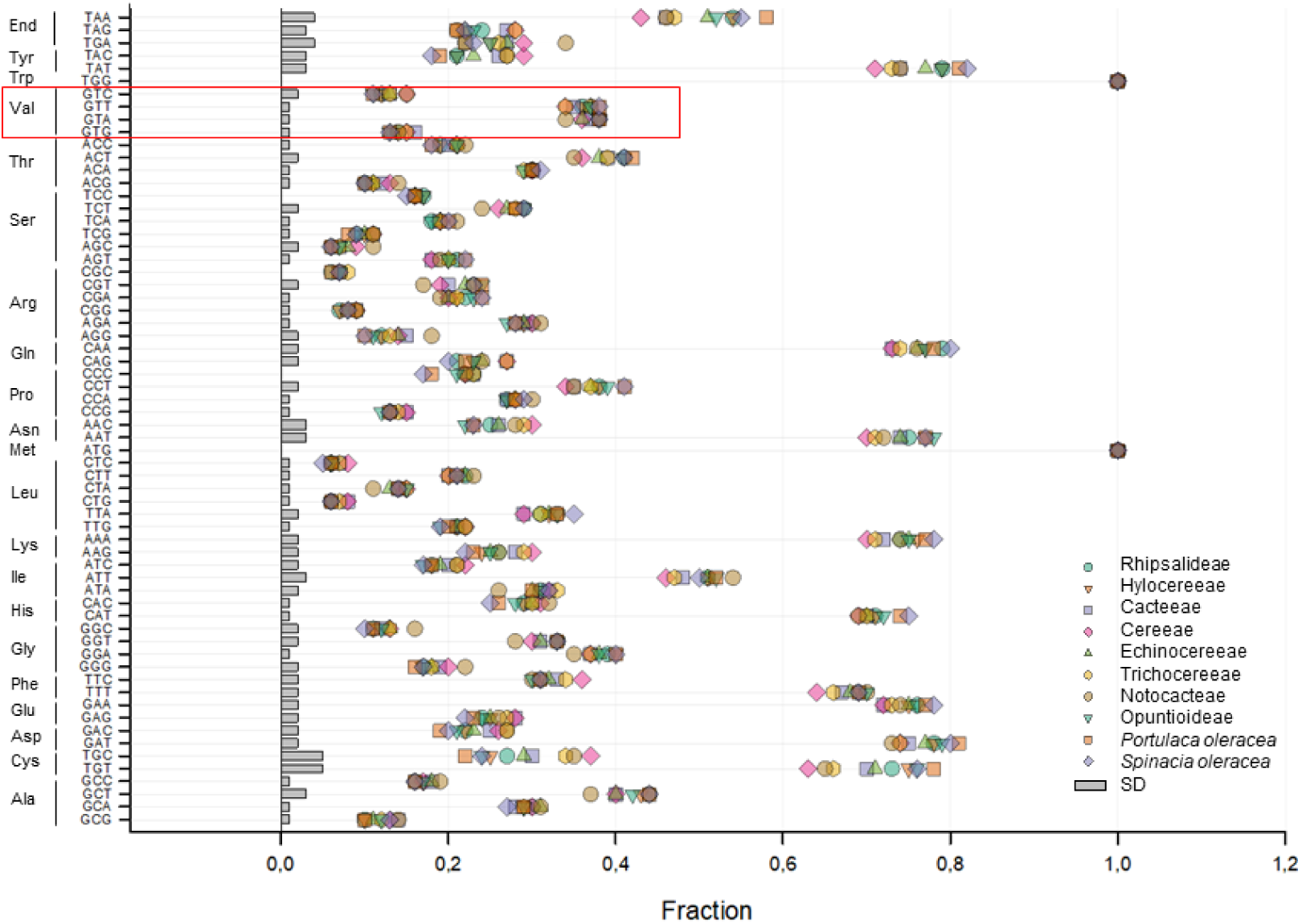
Frequency (colored symbols) and standard deviation (SD, grey bars) of codon usage frequency average of protein-coding genes of Cactoideae tribes, Opuntioideae subfamily, and model species *Portulaca oleracea* and *Spinacea oleracea*. The highlighted section corresponds to the valine codons, whose respective tRNA genes *trnV-GAC* and *trnV-UAC* were lost in both *Cereus* (Tribe Cereeae) analyzed here.

**Supplementary Fig. S4.**
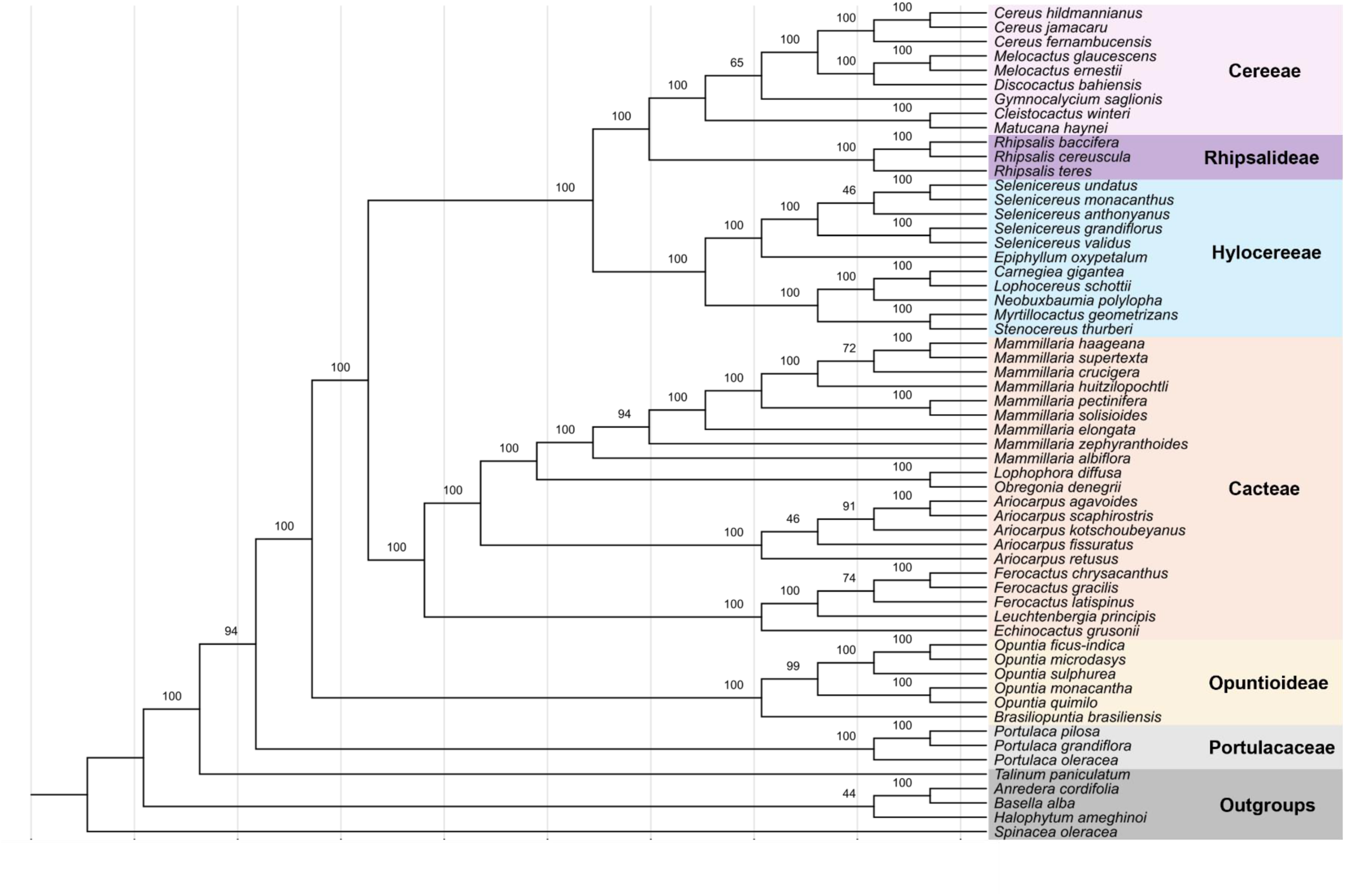
Phylogenetic tree reconstructed by maximum-likelihood based on 45 shared protein-coding genes of 58 cacti species. The bootstrap values are above the node branches. Species are grouped by tribe, indicated in the right. Colored blocks refers to each tribe.

**Supplementary Table S1.**
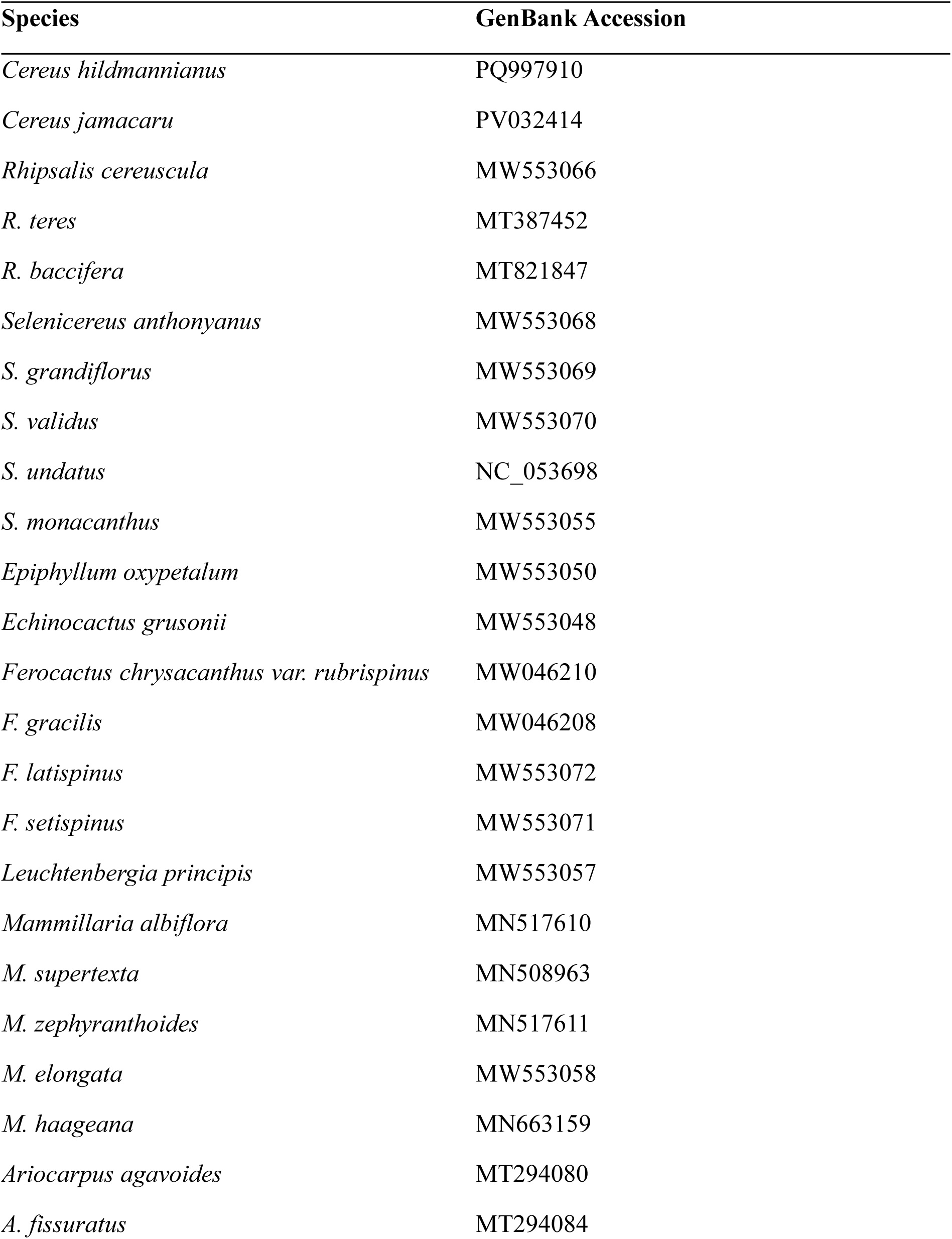
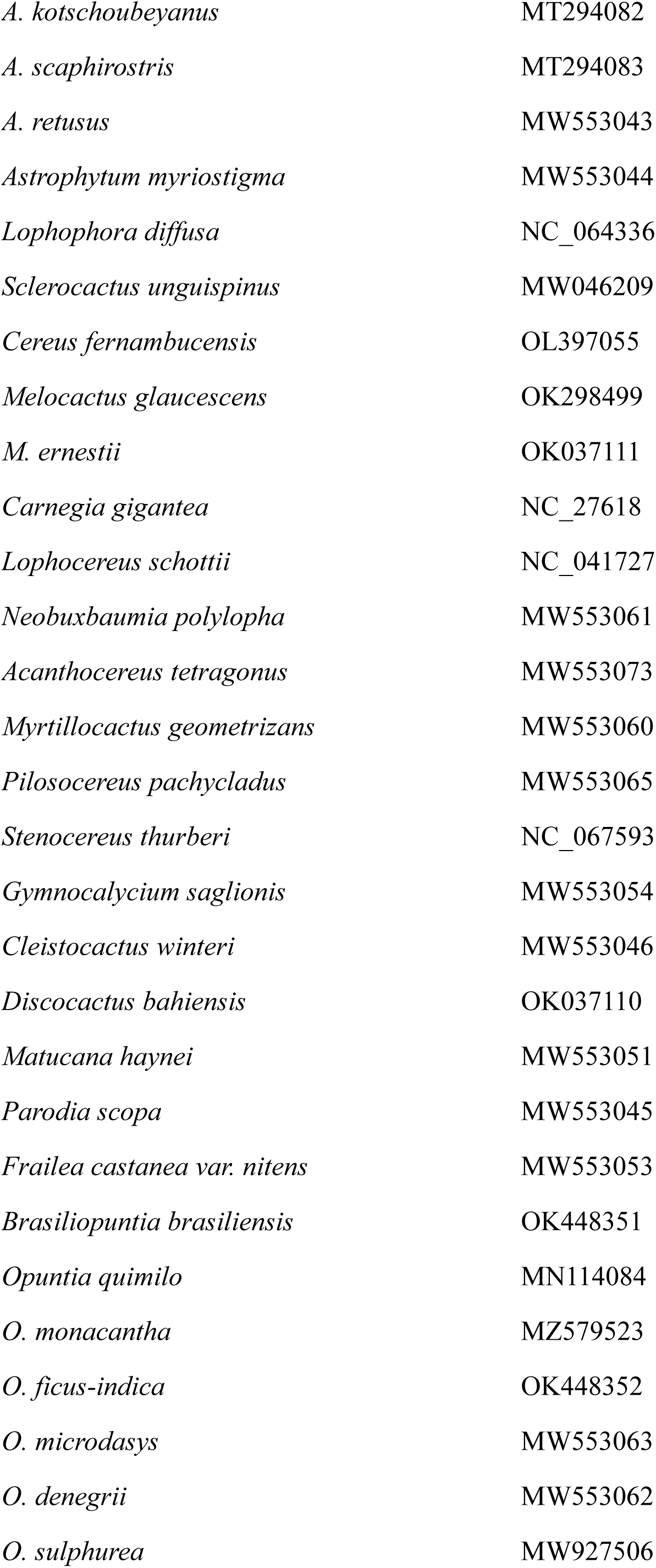
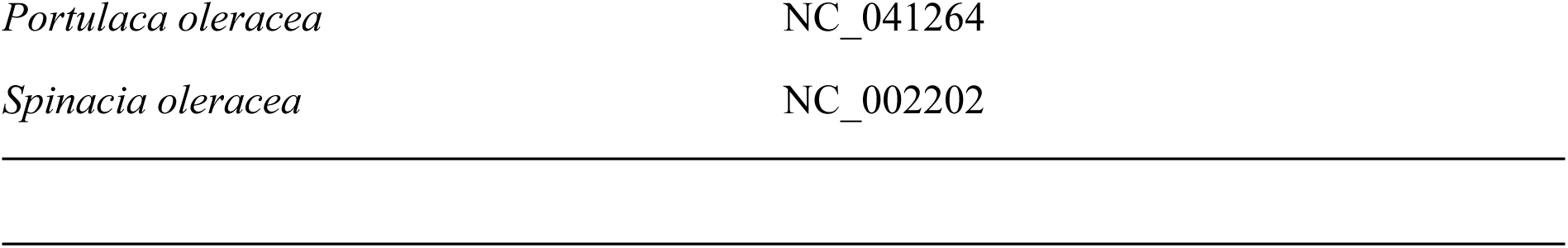
List of species included in the codon usage analysis.

**Supplementary Table S2.** List of species included in the phylogeny.

| Species | Family | GenBank |
| --- | --- | --- |
| *Cereus hildmannianus* | Cactaceae | PQ997910 |
| *Cereus jamacaru* | Cactaceae | PV032414 |
| *Cereus fernambucensis* | Cactaceae | OL397055.1 |
| *Melocactus glaucescens* | Cactaceae | OK298499 |
| *Melocactus ernestii* | Cactaceae | OK037111 |
| *Discocactus bahiensis* | Cactaceae | OK037110 |
| *Gymnocalycium saglionis* | Cactaceae | MW553054 |
| *Cleistocactus winteri* | Cactaceae | MW553046 |
| *Matucana haynei* | Cactaceae | MW553051 |
| *Rhipsalis baccifera* | Cactaceae | MT821847 |
| *Rhipsalis cereuscula* | Cactaceae | MW553066 |
| *Rhipsalis teres* | Cactaceae | MT387452 |
| *Selenicereus undatus* | Cactaceae | NC_053698 |
| *Selenicereus monacanthus* | Cactaceae | MW553055.1 |
| *Selenicereus anthonyanus* | Cactaceae | MW553068.1 |
| *Selenicereus grandiflorus* | Cactaceae | MW553069.1 |
| *Selenicereus validus* | Cactaceae | MW553070.1 |
| *Epiphyllum oxypetalum* | Cactaceae | MW553050 |
| *Carnegiea gigantea* | Cactaceae | NC_027618 |
| *Lophocereus schottii* | Cactaceae | NC_041727 |
| *Neobuxbaumia polylopha* | Cactaceae | MW553061.1 |
| *Myrtillocactus geometrizans* | Cactaceae | MW553060 |
| *Stenocereus thurberi* | Cactaceae | MZ170090.1 |
| *Mammillaria haageana* | Cactaceae | MN663159.1 |
| *Mammillaria supertexta* | Cactaceae | MN508963 |
| *Mammillaria crucigera* | Cactaceae | MN517613.1 |
| *Mammillaria huitzilopochtli* | Cactaceae | MN517612.1 |
| *Mammillaria pectinifera* | Cactaceae | MN519716.1 |
| *Mammillaria solisioides* | Cactaceae | MN518341.1 |
| *Mammillaria elongata* | Cactaceae | MW553058.1 |
| *Mammillaria zephyranthoides* | Cactaceae | MN517611 |
| *Mammillaria albiflora* | Cactaceae | MN517610 |
| *Lophophora diffusa* | Cactaceae | NC_064336.1 |
| *Obregonia denegrii* | Cactaceae | MW553062 |
| *Ariocarpus agavoides* | Cactaceae | MT294080.1 |
| *Ariocarpus scaphiostriis* | Cactaceae | MT294083.1 |
| *Ariocarpus kotschoubeyanus* | Cactaceae | MT294082.1 |
| *Ariocarpus fissuratus* | Cactaceae | MT294084.1 |
| *Ariocarpus retusus* | Cactaceae | MW553043 |
| *Ferocactus chrysacanthus* | Cactaceae | MW046210.1 |
| *Ferocactus gracilis* | Cactaceae | MW046208.1 |
| *Ferocactus latispinus* | Cactaceae | MW553072 |
| *Leuchtenbergia principis* | Cactaceae | MW553057 |
| *Echinocactus grusonii* | Cactaceae | MW553048 |
| *Opuntia ficus-indica* | Cactaceae | OK448352.1 |
| *Opuntia microdasys* | Cactaceae | MW553063 |
| *Opuntia sulphurea* | Cactaceae | MZ491872.1 |
| *Opuntia monacantha* | Cactaceae | MZ579523.1 |
| *Opuntia quimilo* | Cactaceae | MN114084 |
| *Brasiliopuntia brasiliensis* | Cactaceae | OK448351 |
| *Portulaca pilosa* | Portulacaceae | NC_036236.1 |
| *Portulaca grandiflora* | Portulacaceae | NC_041299.1 |
| *Portulaca oleracea* | Portulacaceae | NC_041264.1 |
| *Talinum paniculatum* | Talinaceae | NC_037748.1 |
| *Anredera cordifolia* | Basellaceae | NC_041274.1 |
| *Basella alba* | Basellaceae | NC_041293.1 |
| *Halophytum ameghinoi* | Halophytaceae | NC_040949.1 |
| *Spinacea oleracea* * | Chenopodiaceae* | NC_002202.1 |

\*outgroup

**Supplementary Table S3.**
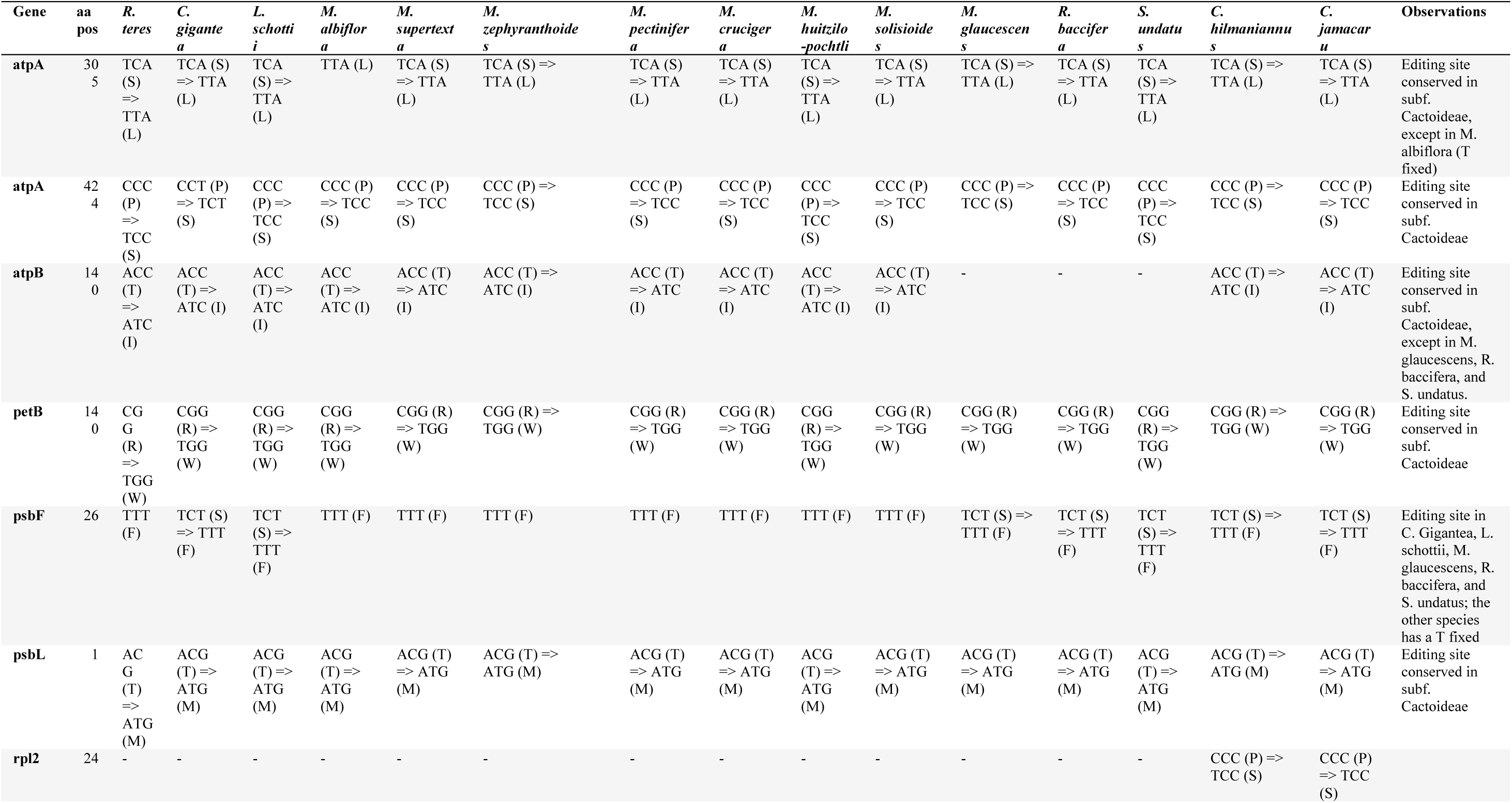
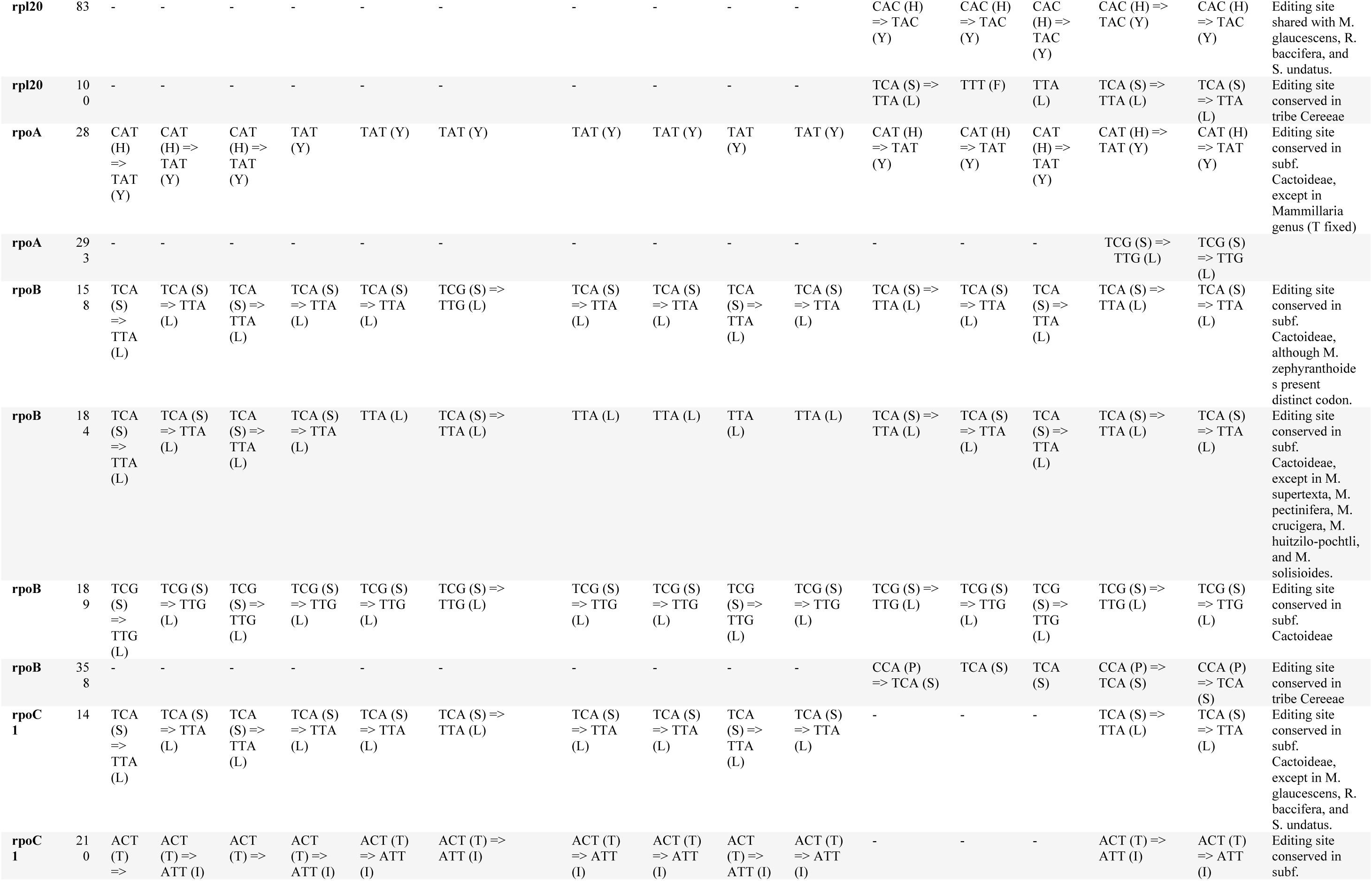
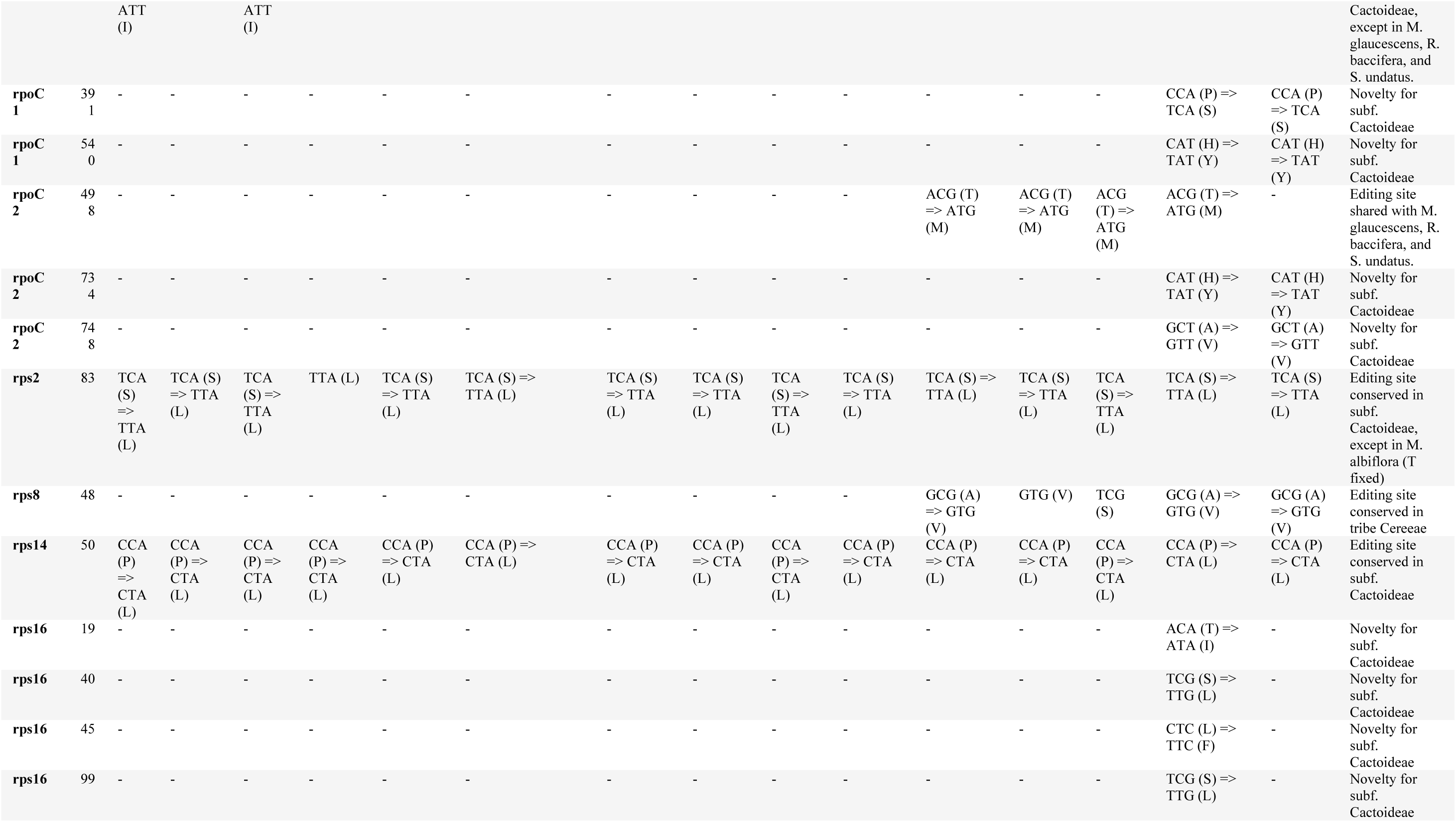
List of RNA editing sites predicted by PREP web server. The abbreviation “aa pos” meaning the amino acid position of the RNA editing site in the aligned sequence.

**Supplementary Table S4.** Distribution of SSR loci in the plastome of *C. hildmannianus* subsp. *hildmannianus*. CDS, coding sequences; IGS, intergenic spacers; LSC, Large Single Copy; IRB, inverted repeat B; SSC, Small Single Copy.

| SSR nr. | SSR type | SSR | Size | Start | End | Location | Region | Genome Region |
| --- | --- | --- | --- | --- | --- | --- | --- | --- |
| 1 | p1 | (T)9 | 9 | 307 | 315 | *ycf3* | intron | LSC |
| 2 | p1 | (A)9 | 9 | 684 | 692 | *ycf3* | intron | LSC |
| 3 | p1 | (A)10 | 10 | 1344 | 1353 | *ycf3* | intron | LSC |
| 4 | p2 | (TA)4 | 8 | 1571 | 1578 | *ycf3* | intron | LSC |
| 5 | p1 | (A)8 | 8 | 1950 | 1957 | *ycf3* | intron | LSC |
| 6 | p1 | (T)11 | 11 | 2373 | 2383 | *ycf3/psaA* | IGS | LSC |
| 7 | c* | (A)8(<A>G)(AG)3 | 16 | 2679 | 2694 | *ycf3/psaA* | IGS | LSC |
| 8 | p2 | (AG)4 | 8 | 7432 | 7439 | *rps14* | CDS | LSC |
| 9 | p2 | (AT)8 | 16 | 8247 | 8262 | *trnG-GCC/psbZ* | IGS | LSC |
| 10 | p2 | (CT)4 | 8 | 9082 | 9089 | *trnS-UGA* | CDS | LSC |
| 11 | p1 | (A)9 | 9 | 9183 | 9191 | *trnS-UGA/psbC* | IGS | LSC |
| 12 | p1 | (A)10 | 10 | 9223 | 9232 | *trnS-UGA/psbC* | IGS | LSC |
| 13 | p1 | (T)9 | 9 | 9247 | 9255 | *trnS-UGA/psbC* | IGS | LSC |
| 14 | p1 | (A)8 | 8 | 12067 | 12074 | *psbD/trnT-GGU* | IGS | LSC |
| 15 | p1 | (A)9 | 9 | 12748 | 12756 | *trnT-GGU/rpoB* | IGS | LSC |
| 16 | p1 | (T)9 | 9 | 12818 | 12826 | *trnT-GGU/rpoB* | IGS | LSC |
| 17 | p1 | (A)10 | 10 | 13664 | 13673 | *rpoB* | CDS | LSC |
| 18 | p1 | (A)8 | 8 | 13770 | 13777 | *rpoB* | CDS | LSC |
| 19 | p2 | (AG)6 | 12 | 15405 | 15416 | *rpoB* | CDS | LSC |
| 20 | p1 | (T)8 | 8 | 16802 | 16809 | *rpoC1* | CDS | LSC |
| 21 | p2 | (AT)5 | 10 | 19240 | 19249 | *rpoC2* | CDS | LSC |
| 22 | p1 | (A)8 | 8 | 20073 | 20080 | *rpoC2* | CDS | LSC |
| 23 | p1 | (T)8 | 8 | 20400 | 20407 | *rpoC2* | CDS | LSC |
| 24 | p1 | (A)11 | 11 | 20541 | 20551 | *rpoC2* | CDS | LSC |
| 25 | p2 | (AT)4 | 8 | 20578 | 20585 | *rpoC2* | CDS | LSC |
| 26 | p1 | (A)8 | 8 | 20712 | 20719 | *rpoC2* | CDS | LSC |
| 27 | p1 | (T)8 | 8 | 21362 | 21369 | *rpoC2* | CDS | LSC |
| 28 | p1 | (A)9 | 9 | 22424 | 22432 | *rpoC2* | CDS | LSC |
| 29 | p1 | (A)9 | 9 | 22594 | 22602 | *rpoC2/rps2* | IGS | LSC |
| 30 | p1 | (A)8 | 8 | 23238 | 23245 | *rps2* | CDS | LSC |
| 31 | p1 | (T)8 | 8 | 24852 | 24859 | *atpI/atpH* | IGS | LSC |
| 32 | p1 | (T)8 | 8 | 25637 | 25644 | *atpF* | intron | LSC |
| 33 | p1 | (A)8 | 8 | 25661 | 25668 | *atpF* | intron | LSC |
| 34 | p1 | (A)9 | 9 | 25874 | 25882 | *atpF* | intron | LSC |
| 35 | p1 | (T)8 | 8 | 25962 | 25969 | *atpF* | intron | LSC |
| 36 | p1 | (A)9 | 9 | 26052 | 26060 | *atpF* | intron | LSC |
| 37 | p1 | (T)10 | 10 | 26130 | 26139 | *atpF* | intron | LSC |
| 38 | p1 | (T)8 | 8 | 26151 | 26158 | *atpF* | intron | LSC |
| 39 | p1 | (T)12 | 12 | 26769 | 26780 | *atpF/atpA* | IGS | LSC |
| 40 | p1 | (A)11 | 11 | 28420 | 28430 | *atpA/trnR-UCU* | IGS | LSC |
| 41 | p1 | (A)9 | 9 | 29060 | 29068 | *trnG-UCC* | intron | LSC |
| 42 | p1 | (A)9 | 9 | 29137 | 29145 | *trnG-UCC* | intron | LSC |
| 43 | p1 | (T)8 | 8 | 29452 | 29459 | *trnG-UCC/clpP* | IGS | LSC |
| 44 | p1 | (A)8 | 8 | 29947 | 29954 | *clpP* | CDS | LSC |
| 45 | p2 | (AT)4 | 8 | 30411 | 30418 | *clpP/trnS-GCU* | IGS | LSC |
| 46 | p2 | (TA)6 | 12 | 30619 | 30630 | *trnS-GCU/psbI* | IGS | LSC |
| 47 | p1 | (A)8 | 8 | 30816 | 30823 | *psbI/psbK* | IGS | LSC |
| 48 | p1 | (T)8 | 8 | 31348 | 31355 | *psbK/trnQ-UUG* | IGS | LSC |
| 49 | p1 | (A)8 | 8 | 31903 | 31910 | *trnQ-UUG/cemA* | IGS | LSC |
| 50 | p1 | (T)8 | 8 | 31964 | 31971 | *trnQ-UUG/cemA* | IGS | LSC |
| 51 | p3 | (AAT)4 | 12 | 32023 | 32034 | *trnQ-UUG/cemA* | IGS | LSC |
| 52 | p1 | (A)11 | 11 | 32036 | 32046 | *cemA* | CDS | LSC |
| 53 | p2 | (AT)4 | 8 | 33056 | 33063 | *petA* | CDS | LSC |
| 54 | p1 | (A)8 | 8 | 33416 | 33423 | *petA* | CDS | LSC |
| 55 | p1 | (T)10 | 10 | 33991 | 34000 | *petA/psbJ* | IGS | LSC |
| 56 | p1 | (T)9 | 9 | 34016 | 34024 | *petA/psbJ* | IGS | LSC |
| 57 | p1 | (T)9 | 9 | 34034 | 34042 | *petA/psbJ* | IGS | LSC |
| 58 | p1 | (T)9 | 9 | 35768 | 35776 | *petL/petG* | IGS | LSC |
| 59 | p1 | (A)8 | 8 | 36515 | 36522 | *trnP-UGG/psaJ* | IGS | LSC |
| 60 | p1 | (T)11 | 11 | 37045 | 37055 | *rpl33/rps18* | IGS | LSC |
| 61 | p2 | (TA)11 | 22 | 37066 | 37087 | *rpl33/rps18* | IGS | LSC |
| 62 | p1 | (A)9 | 9 | 37090 | 37098 | *rpl33/rps18* | IGS | LSC |
| 63 | p1 | (A)8 | 8 | 37423 | 37430 | *rps18* | CDS | LSC |
| 64 | p1 | (A)8 | 8 | 37753 | 37760 | *rps18* | CDS | LSC |
| 65 | p1 | (T)8 | 8 | 38541 | 38548 | *rpl20* | IGS | LSC |
| 66 | p1 | (A)10 | 10 | 38820 | 38829 | *rpl20/rps12* | IGS | LSC |
| 67 | p1 | (A)8 | 8 | 39835 | 39842 | *clpP-fragment* | IGS | LSC |
| 68 | p1 | (T)8 | 8 | 41989 | 41996 | *psbB/psbT* | IGS | LSC |
| 69 | p1 | (T)9 | 9 | 43131 | 43139 | *petB* | intron | LSC |
| 70 | p1 | (A)8 | 8 | 43334 | 43341 | *petB* | intron | LSC |
| 71 | p1 | (A)11 | 11 | 44460 | 44470 | *petB/petD* | IGS | LSC |
| 72 | p2 | (AG)4 | 8 | 44601 | 44608 | *petD* | intron | LSC |
| 73 | p1 | (T)8 | 8 | 44723 | 44730 | *petD* | intron | LSC |
| 74 | p2 | (AT)4 | 8 | 44879 | 44886 | *petD* | intron | LSC |
| 75 | p2 | (AT)6 | 12 | 44889 | 44900 | *petD* | intron | LSC |
| 76 | p1 | (T)15 | 15 | 46090 | 46104 | *rpoA* | CDS | LSC |
| 77 | p2 | (GA)4 | 8 | 46776 | 46783 | *rpoA* | CDS | LSC |
| 78 | p1 | (T)8 | 8 | 46906 | 46913 | *rpoA/rps11* | IGS | LSC |
| 79 | p1 | (T)9 | 9 | 47427 | 47435 | *rps11/rpl36* | IGS | LSC |
| 80 | p1 | (T)12 | 12 | 48518 | 48529 | *rps8/rpl14* | IGS | LSC |
| 81 | p1 | (A)9 | 9 | 48544 | 48552 | *rps8/rpl14* | IGS | LSC |
| 82 | p5 | (AAAGG)3 | 15 | 48557 | 48571 | *rps8/rpl14* | IGS | LSC |
| 83 | p4 | (TTTC)3 | 12 | 50135 | 50146 | *rpl16* | intron | LSC |
| 84 | p1 | (A)9 | 9 | 50226 | 50234 | *rpl16* | intron | LSC |
| 85 | p1 | (T)9 | 9 | 51326 | 51334 | *rpl22* | CDS | LSC |
| 86 | p1 | (T)11 | 11 | 52197 | 52207 | *rps19/rpl2* | IGS | LSC |
| 87 | p4 | (ATCA)3 | 12 | 53570 | 53581 | *rps2/trnS-GGA* | IGS | IRB |
| 88 | p2 | (TA)5 | 10 | 53924 | 53933 | *trnS-GGA/rps4* | IGS | IRB |
| 89 | p1 | (T)8 | 8 | 54925 | 54932 | *rps4/trnT-UGU* | IGS | IRB |
| 90 | p1 | (A)8 | 8 | 55015 | 55022 | *rps4/trnT-UGU* | IGS | IRB |
| 91 | p1 | (A)8 | 8 | 55035 | 55042 | *rps4/trnT-UGU* | IGS | IRB |
| 92 | p1 | (A)8 | 8 | 55529 | 55536 | *trnL-UAA* | intron | IRB |
| 93 | p1 | (A)9 | 9 | 55567 | 55575 | *trnL-UAA* | intron | IRB |
| 94 | p1 | (A)8 | 8 | 55669 | 55676 | *trnL-UAA* | intron | IRB |
| 95 | p2 | (TA)7 | 14 | 55743 | 55756 | *trnL-UAA* | intron | IRB |
| 96 | p2 | (TA)5 | 10 | 55761 | 55770 | *trnL-UAA* | intron | IRB |
| 97 | p2 | (AT)4 | 8 | 55813 | 55820 | *trnL-UAA* | intron | IRB |
| 98 | p1 | (T)8 | 8 | 56163 | 56170 | *trnL-UAA/trnF-GAA* | IGS | IRB |
| 99 | p1 | (C)12 | 12 | 57309 | 57320 | *trnF-GAA/ndhJ* | IGS | IRB |
| 100 | p1 | (G)9 | 9 | 57323 | 57331 | *trnF-GAA/ndhJ* | IGS | IRB |
| 101 | p1 | (A)8 | 8 | 57938 | 57945 | *ndhJ/trnD-GUC* | IGS | IRB |
| 102 | p1 | (T)8 | 8 | 58271 | 58278 | *trnD-GUC/psbM* | IGS | IRB |
| 103 | p2 | (TA)7 | 14 | 58673 | 58686 | *trnD-GUC/psbM* | IGS | IRB |
| 104 | p1 | (T)8 | 8 | 59308 | 59315 | *psbM/petN* | IGS | IRB |
| 105 | p1 | (A)8 | 8 | 59499 | 59506 | *psbM/petN* | IGS | IRB |
| 106 | p1 | (A)9 | 9 | 60320 | 60328 | *petN/trnC-GCA* | IGS | IRB |
| 107 | p1 | (T)8 | 8 | 60761 | 60768 | *petN/trnC-GCA* | IGS | IRB |
| 108 | p1 | (T)8 | 8 | 60895 | 60902 | *trnC-GCA/trnE-UUC* | IGS | IRB |
| 109 | p2 | (AT)6 | 12 | 60993 | 61004 | *trnC-GCA/trnE-UUC* | IGS | IRB |
| 110 | p1 | (A)11 | 11 | 61217 | 61227 | *trnC-GCA/trnE-UUC* | IGS | IRB |
| 111 | p1 | (T)8 | 8 | 61736 | 61743 | *trnY-GUA/ndhC* | IGS | IRB |
| 112 | p1 | (A)8 | 8 | 61775 | 61782 | *trnY-GUA/ndhC* | IGS | IRB |
| 113 | p2 | (AT)8 | 16 | 61997 | 62012 | *ndhC/rbcL* | IGS | IRB |
| 114 | p2 | (TC)4 | 8 | 63128 | 63135 | *rbcL* | CDS | IRB |
| 115 | p2 | (AT)4 | 8 | 63803 | 63810 | *rbcL/atpB* | IGS | IRB |
| 116 | p5 | (TTTAT)3 | 15 | 64316 | 64330 | *rbcL/atpB* | IGS | IRB |
| 117 | p1 | (A)10 | 10 | 64431 | 64440 | *atpB* | CDS | IRB |
| 118 | p2 | (TA)4 | 8 | 66364 | 66371 | *atpE/trnM-CAU* | IGS | IRB |
| 119 | p1 | (T)8 | 8 | 67087 | 67094 | *trnM-CAU/accD* | IGS | IRB |
| 120 | p3 | (GTT)4 | 12 | 68674 | 68685 | *accD* | CDS | IRB |
| 121 | p2 | (TA)4 | 8 | 69236 | 69243 | *accD* | CDS | IRB |
| 122 | p4 | (ATAA)3 | 12 | 69539 | 69550 | *accD/psaI* | IGS | IRB |
| 123 | p4 | (AATT)3 | 12 | 69634 | 69645 | *accD/psaI* | IGS | IRB |
| 124 | p1 | (A)8 | 8 | 69657 | 69664 | *accD/psaI* | IGS | IRB |
| 125 | p1 | (G)8 | 8 | 69961 | 69968 | *psaI/ycf4* | IGS | IRB |
| 126 | p1 | (C)8 | 8 | 70280 | 70287 | *ycf4* | CDS | IRB |
| 127 | p1 | (T)11 | 11 | 70294 | 70304 | *ycf4* | CDS | IRB |
| 128 | p1 | (T)8 | 8 | 70734 | 70741 | *ycf4/rps16* | IGS | IRB |
| 129 | p2 | (AT)7 | 14 | 70778 | 70791 | *ycf4/rps16* | IGS | IRB |
| 130 | p2 | (TA)5 | 10 | 71269 | 71278 | *rps16* | intron | IRB |
| 131 | p1 | (A)8 | 8 | 71402 | 71409 | *rps16* | intron | IRB |
| 132 | p1 | (T)8 | 8 | 71572 | 71579 | *rps16* | intron | IRB |
| 133 | p1 | (A)10 | 10 | 72032 | 72041 | *rps16/trnK-UUU* | IGS | IRB |
| 134 | p1 | (A)9 | 9 | 72462 | 72470 | *trnK-UUU* | intron | IRB |
| 135 | p1 | (T)8 | 8 | 72817 | 72824 | *trnK-UUU* | intron | IRB |
| 136 | p1 | (A)9 | 9 | 73200 | 73208 | *trnK-UUU* | intron | IRB |
| 137 | p1 | (T)8 | 8 | 73846 | 73853 | *trnK-UUU* | intron | IRB |
| 138 | p1 | (A)10 | 10 | 74430 | 74439 | *trnK-UUU* | intron | IRB |
| 139 | p1 | (T)10 | 10 | 76085 | 76094 | *psbA/trnH-GUG* | IGS | IRB |
| 140 | p1 | (T)9 | 9 | 76414 | 76422 | *trnH-GUG/trnI-CAU* | IGS | IRB |
| 141 | p4 | (ATAA)3 | 12 | 76765 | 76776 | *trnI-CAU/ycf2* | IGS | IRB |
| 142 | p2 | (GA)4 | 8 | 76873 | 76880 | *ycf2* | CDS | IRB |
| 143 | p2 | (GA)4 | 8 | 76885 | 76892 | *ycf2* | CDS | IRB |
| 144 | p2 | (GA)4 | 8 | 77818 | 77825 | *ycf2* | CDS | IRB |
| 145 | p1 | (A)9 | 9 | 79793 | 79801 | *ycf2* | CDS | IRB |
| 146 | p2 | (GA)4 | 8 | 79814 | 79821 | *ycf2* | CDS | IRB |
| 147 | p3 | (GAA)4 | 12 | 80354 | 80365 | *ycf2* | CDS | IRB |
| 148 | p1 | (A)11 | 11 | 84593 | 84603 | *ycf1* | CDS | IRB |
| 149 | p3 | (GAA)9 | 27 | 84625 | 84651 | *ycf1* | CDS | IRB |
| 150 | p1 | (A)9 | 9 | 85151 | 85159 | *ycf1* | CDS | IRB |
| 151 | p1 | (A)11 | 11 | 85184 | 85194 | *ycf1* | CDS | IRB |
| 152 | p1 | (A)8 | 8 | 85311 | 85318 | *ycf1* | CDS | SSC |
| 153 | p1 | (A)9 | 9 | 85395 | 85403 | *ycf1* | CDS | SSC |
| 154 | p1 | (T)8 | 8 | 86603 | 86610 | *ycf1* | CDS | SSC |
| 155 | p1 | (A)8 | 8 | 87090 | 87097 | *ycf1* | CDS | SSC |
| 156 | p1 | (A)8 | 8 | 87540 | 87547 | *ycf1* | CDS | SSC |
| 157 | p1 | (A)9 | 9 | 87574 | 87582 | *ycf1* | CDS | SSC |
| 158 | p4 | (AATT)3 | 12 | 87605 | 87616 | *ycf1* | CDS | SSC |
| 159 | p1 | (A)8 | 8 | 87661 | 87668 | *ycf1* | CDS | SSC |
| 160 | p1 | (A)9 | 9 | 88257 | 88265 | *ycf1* | CDS | SSC |
| 161 | p1 | (A)8 | 8 | 88267 | 88274 | *ycf1* | CDS | SSC |
| 162 | p6 | (AAGAAA)4 | 24 | 88418 | 88441 | *ycf1* | CDS | SSC |
| 163 | p1 | (A)10 | 10 | 88511 | 88520 | *ycf1* | CDS | SSC |
| 164 | p1 | (A)9 | 9 | 89564 | 89572 | *rps15* | CDS | SSC |
| 165 | p4 | (CCAT)3 | 12 | 89988 | 89999 | *rps15/ndhH* | IGS | SSC |
| 166 | p1 | (A)9 | 9 | 90781 | 90789 | *ndhH/trnL-UAG* | IGS | SSC |
| 167 | p1 | (T)8 | 8 | 91082 | 91089 | *trnL-UAG/ccsA* | IGS | SSC |
| 168 | p1 | (T)8 | 8 | 92266 | 92273 | *ccsA/ndhD* | IGS | SSC |
| 169 | p1 | (A)8 | 8 | 92305 | 92312 | *ccsA/ndhD* | IGS | SSC |
| 170 | p1 | (A)9 | 9 | 92435 | 92443 | *ccsA/ndhD* | IGS | SSC |
| 171 | p1 | (A)11 | 11 | 93525 | 93535 | *ndhD/psaC* | IGS | SSC |
| 172 | p1 | (A)10 | 10 | 93846 | 93855 | *ndhD/psaC* | IGS | SSC |
| 173 | p1 | (T)8 | 8 | 94158 | 94165 | *psaC/ndhH-fragment* | IGS | SSC |
| 174 | p5 | (AGTAA)3 | 15 | 94193 | 94207 | *psaC/ndhH-fragment* | IGS | SSC |
| 175 | p1 | (T)8 | 8 | 94500 | 94507 | *psaC/ndhH-fragment* | IGS | SSC |
| 176 | p1 | (A)8 | 8 | 95089 | 95096 | *ndhH-fragment/rpl32* | IGS | SSC |
| 177 | p1 | (A)11 | 11 | 95370 | 95380 | *ndhH-fragment/rpl32* | IGS | SSC |
| 178 | p1 | (T)10 | 10 | 95402 | 95411 | *ndhH-fragment/rpl32* | IGS | SSC |
| 179 | p1 | (T)14 | 14 | 95534 | 95547 | *rpl32* | CDS | SSC |
| 180 | p1 | (T)8 | 8 | 95686 | 95693 | *rpl32* | CDS | SSC |
| 181 | p2 | (AT)7 | 14 | 95917 | 95930 | *rpl32/trnN-GUU* | IGS | SSC |
| 182 | p1 | (T)10 | 10 | 96135 | 96144 | *rpl32/trnN-GUU* | IGS | SSC |
| 183 | p1 | (A)9 | 9 | 96609 | 96617 | *ndhF* | CDS | SSC |
| **184** | p2 | (CA)4 | 8 | 97236 | 97243 | *ndhF/trnR-ACG* | IGS | SSC |
| **185** | p1 | (A)9 | 9 | 97449 | 97457 | *trnR-ACG/rrn5* | IGS | SSC |
| **186** | p2 | (GA)4 | 8 | 97766 | 97773 | *rrn5/rrn4.5* | IGS | SSC |
| **187** | p2 | (AG)4 | 8 | 99209 | 99216 | *rrn23* | CDS | SSC |
| **188** | p1 | (A)9 | 9 | 102585 | 102593 | *trnI-GAU* | intron | SSC |
| **189** | p1 | (T)8 | 8 | 105724 | 105731 | *rrn16/rps12* | IGS | SSC |
| **190** | p2 | (AT)5 | 10 | 105786 | 105795 | *rrn16/rps12* | IGS | SSC |
| **191** | p2 | (TC)4 | 8 | 109537 | 109544 | *ndhB/ycf1* | IGS | SSC |

**Supplementary Table S5.**
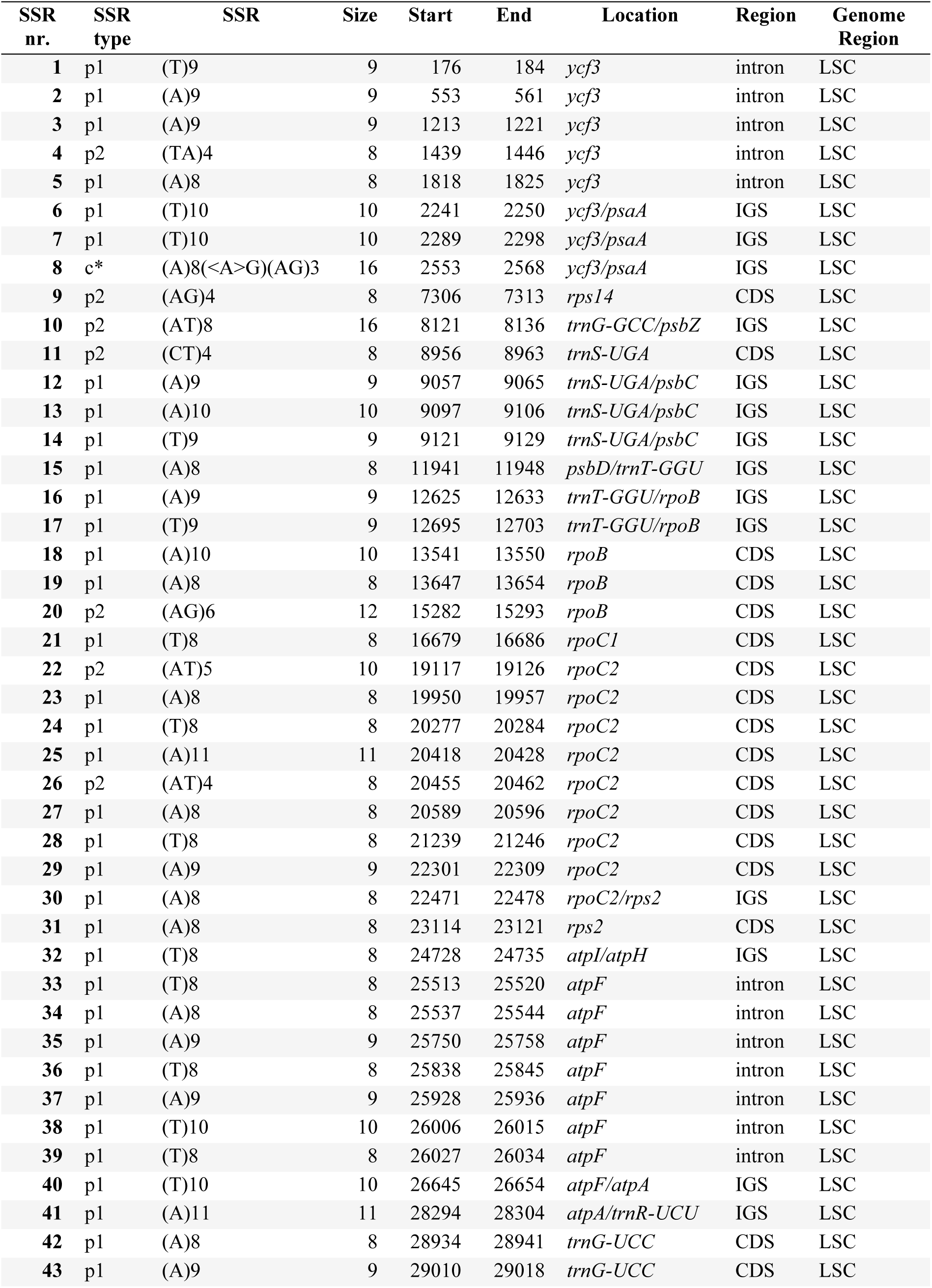
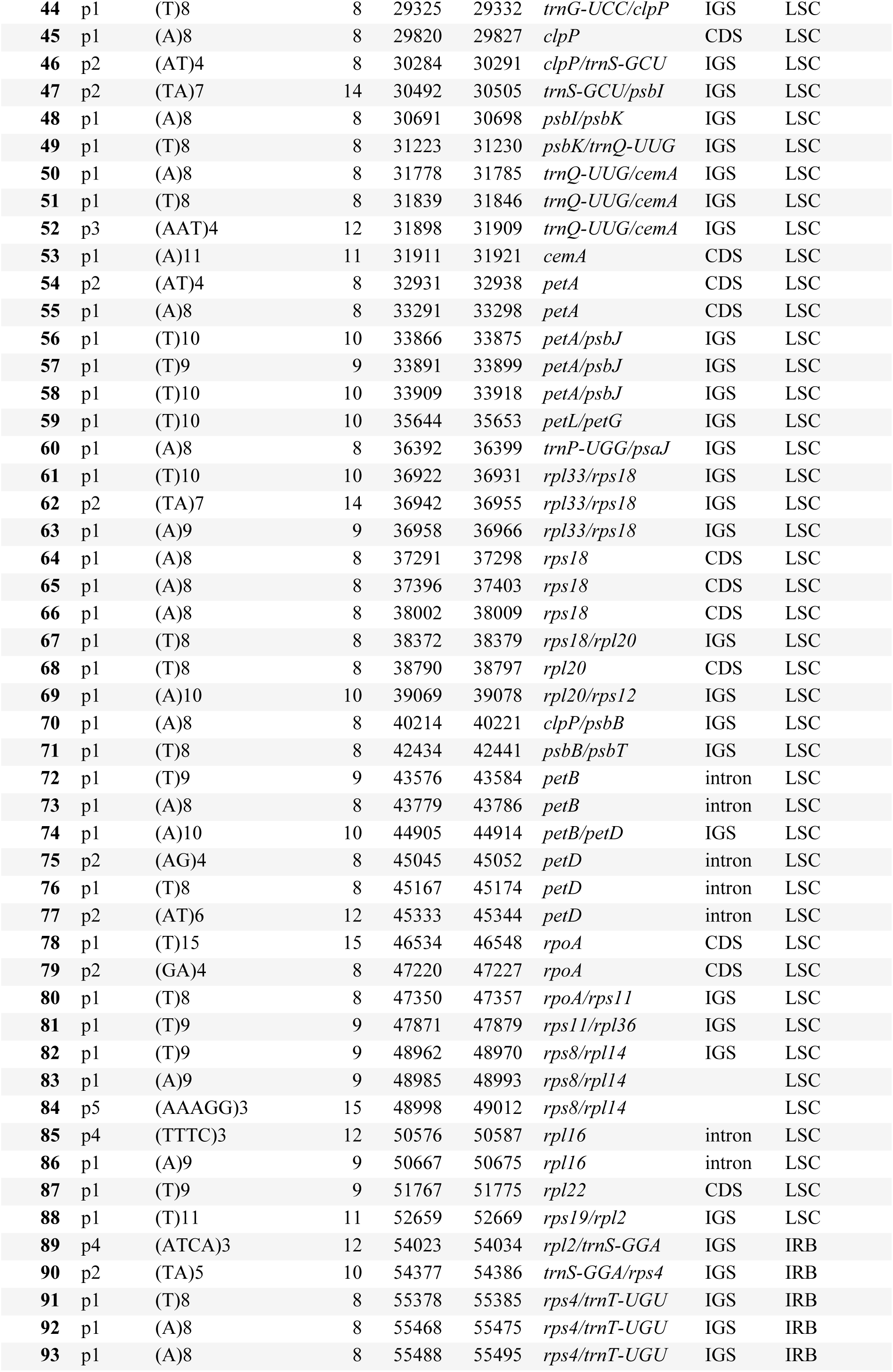
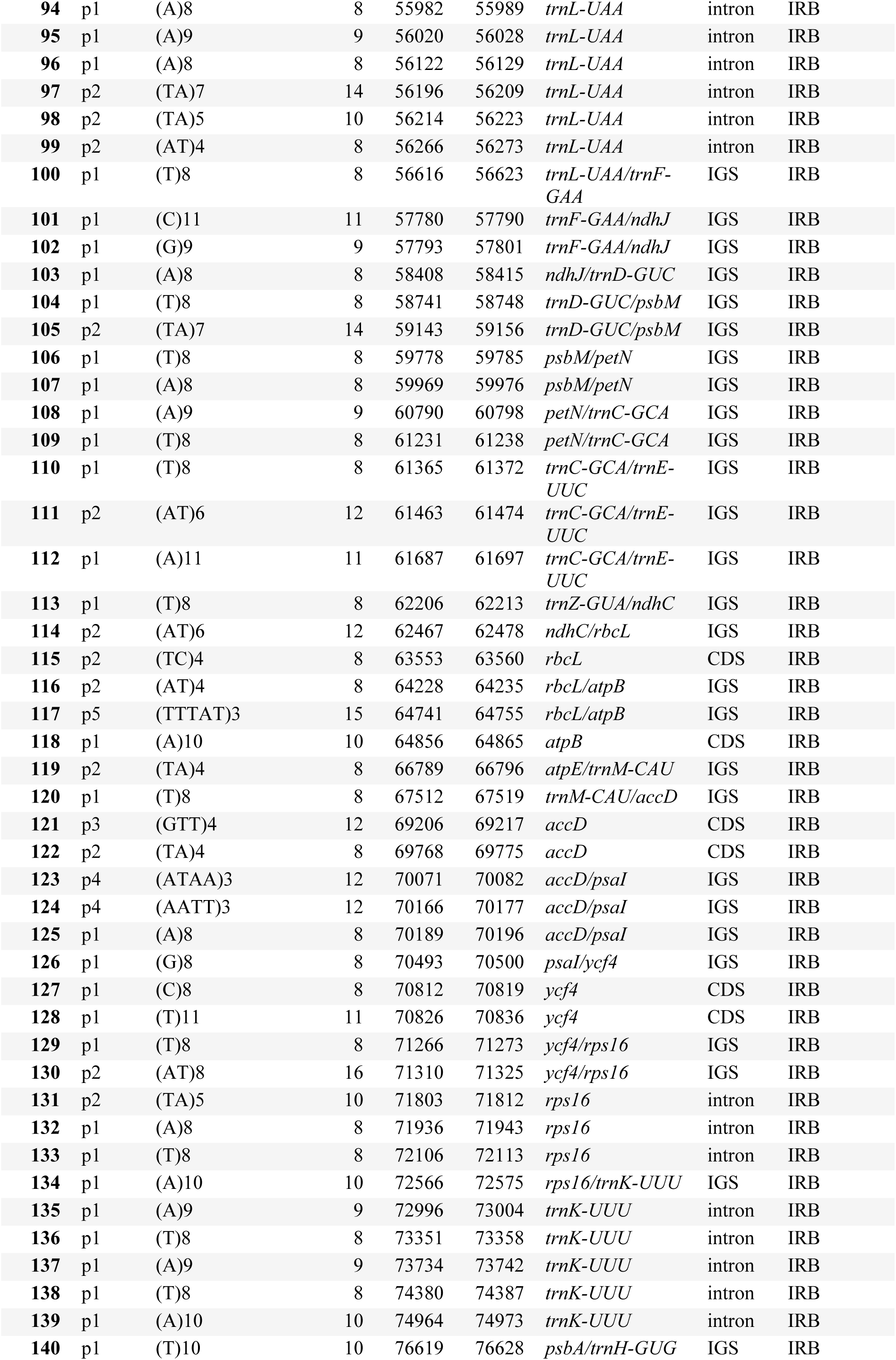
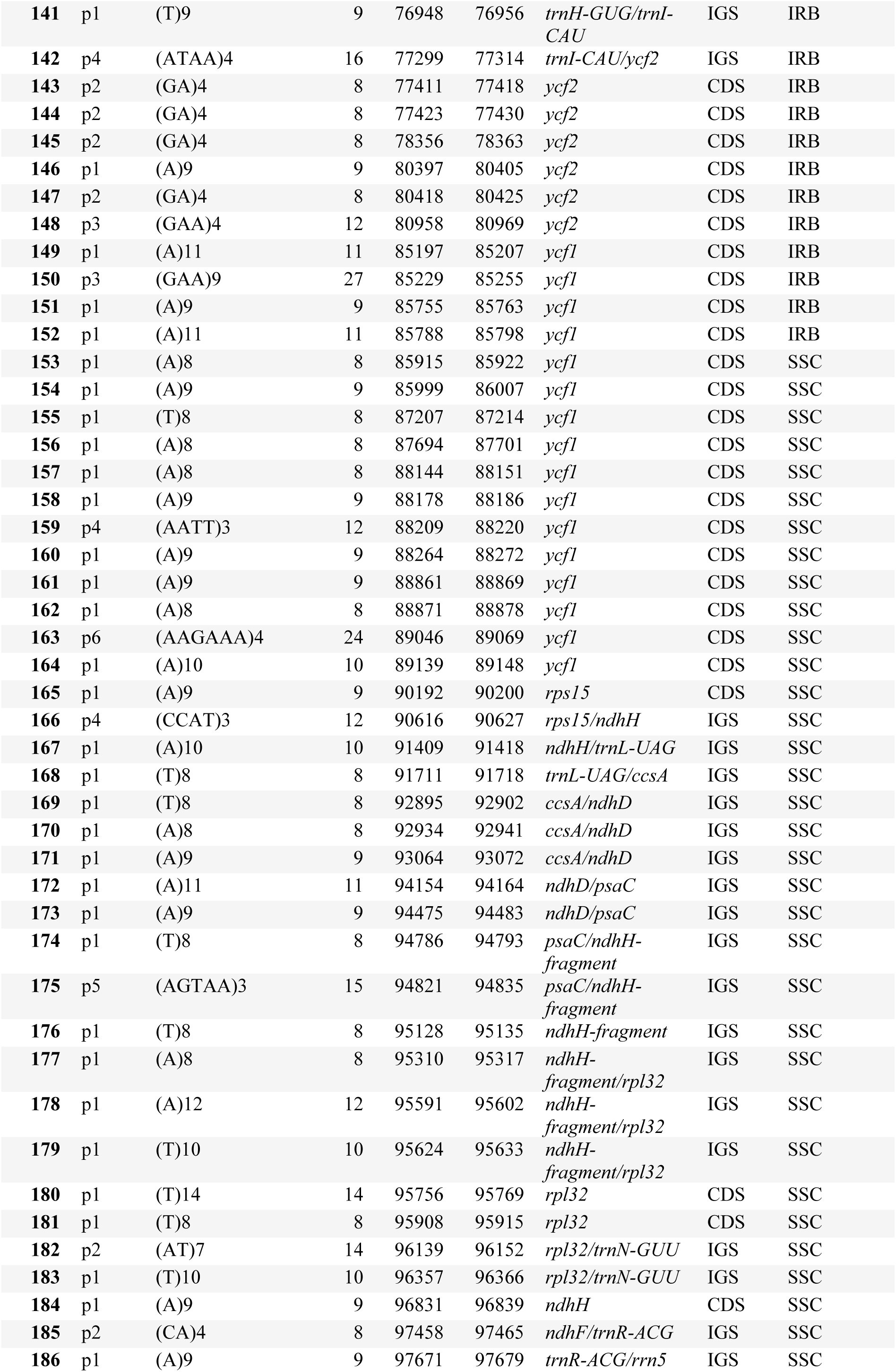
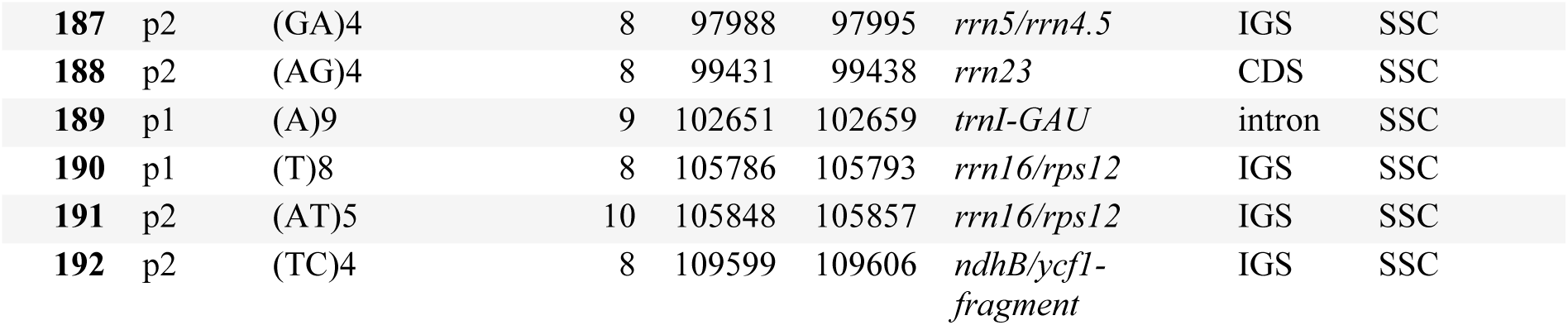
Distribution of SSR loci in the plastome of *C. jamacaru* subsp. *jamacaru*. CDS, coding sequences; IGS, intergenic spacers; LSC, Large Single Copy; IRB, inverted repeat B; SSC, Small Single Copy.

**Supplementary Table S6.**
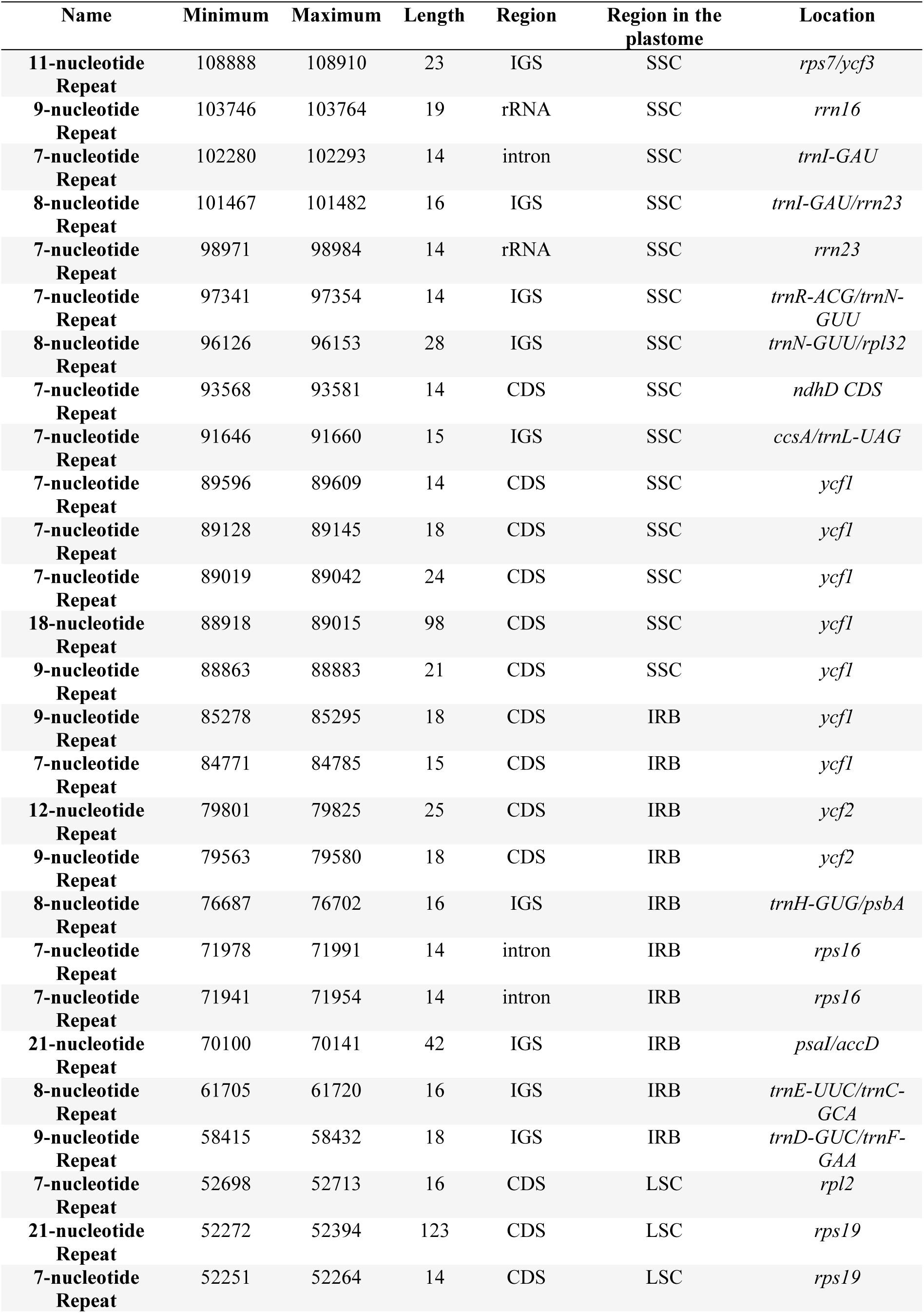
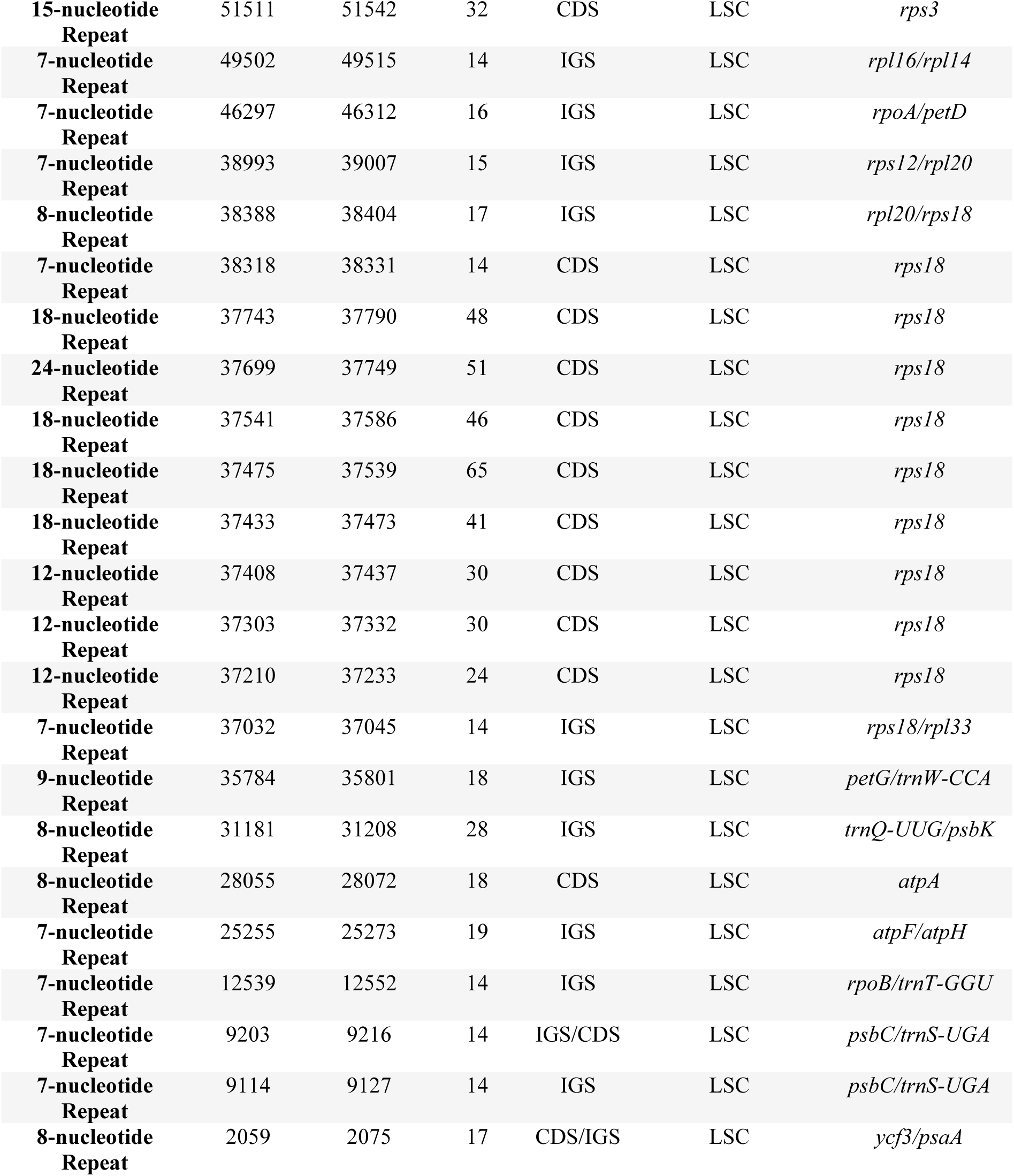
Distribution of tandem repeats in the plastome of *C. jamacaru* subsp. *jamacaru*. CDS, coding sequences; IGS, intergenic spacers.

| Name | Minimum | Maximum | Length | Region | Region in the plastome | Location |
| --- | --- | --- | --- | --- | --- | --- |
| 11-nucleotide Repeat | 108888 | 108910 | 23 | IGS | SSC | *rps7/ycf3* |
| 9-nucleotide Repeat | 103746 | 103764 | 19 | rRNA | SSC | *rrn16* |
| 7-nucleotide Repeat | 102280 | 102293 | 14 | intron | SSC | *trnI-GAU* |
| 8-nucleotide Repeat | 101467 | 101482 | 16 | IGS | SSC | *trnI-GAU/rrn23* |
| 7-nucleotide Repeat | 98971 | 98984 | 14 | rRNA | SSC | *rrn23* |
| 7-nucleotide Repeat | 97341 | 97354 | 14 | IGS | SSC | *trnR-ACG/trnN-GUU* |
| 8-nucleotide Repeat | 96126 | 96153 | 28 | IGS | SSC | *trnN-GUU/rpl32* |
| 7-nucleotide Repeat | 93568 | 93581 | 14 | CDS | SSC | *ndhD CDS* |
| 7-nucleotide Repeat | 91646 | 91660 | 15 | IGS | SSC | *ccsA/trnL-UAG* |
| 7-nucleotide Repeat | 89596 | 89609 | 14 | CDS | SSC | *ycf1* |
| 7-nucleotide Repeat | 89128 | 89145 | 18 | CDS | SSC | *ycf1* |
| 7-nucleotide Repeat | 89019 | 89042 | 24 | CDS | SSC | *ycf1* |
| 18-nucleotide Repeat | 88918 | 89015 | 98 | CDS | SSC | *ycf1* |
| 9-nucleotide Repeat | 88863 | 88883 | 21 | CDS | SSC | *ycf1* |
| 9-nucleotide Repeat | 85278 | 85295 | 18 | CDS | IRB | *ycf1* |
| 7-nucleotide Repeat | 84771 | 84785 | 15 | CDS | IRB | *ycf1* |
| 12-nucleotide Repeat | 79801 | 79825 | 25 | CDS | IRB | *ycf2* |
| 9-nucleotide Repeat | 79563 | 79580 | 18 | CDS | IRB | *ycf2* |
| 8-nucleotide Repeat | 76687 | 76702 | 16 | IGS | IRB | *trnH-GUG/psbA* |
| 7-nucleotide Repeat | 71978 | 71991 | 14 | intron | IRB | *rps16* |
| 7-nucleotide Repeat | 71941 | 71954 | 14 | intron | IRB | *rps16* |
| 21-nucleotide Repeat | 70100 | 70141 | 42 | IGS | IRB | *psaI/accD* |
| 8-nucleotide Repeat | 61705 | 61720 | 16 | IGS | IRB | *trnE-UUC/trnC-GCA* |
| 9-nucleotide Repeat | 58415 | 58432 | 18 | IGS | IRB | *trnD-GUC/trnF-GAA* |
| 7-nucleotide Repeat | 52698 | 52713 | 16 | CDS | LSC | *rpl2* |
| 21-nucleotide Repeat | 52272 | 52394 | 123 | CDS | LSC | *rps19* |
| 7-nucleotide Repeat | 52251 | 52264 | 14 | CDS | LSC | *rps19* |
| **15-nucleotide Repeat** | 51511 | 51542 | 32 | CDS | LSC | *rps3* |
| **7-nucleotide Repeat** | 49502 | 49515 | 14 | IGS | LSC | *rpl16/rpl14* |
| **7-nucleotide Repeat** | 46297 | 46312 | 16 | IGS | LSC | *rpoA/petD* |
| **7-nucleotide Repeat** | 38993 | 39007 | 15 | IGS | LSC | *rps12/rpl20* |
| **8-nucleotide Repeat** | 38388 | 38404 | 17 | IGS | LSC | *rpl20/rps18* |
| **7-nucleotide Repeat** | 38318 | 38331 | 14 | CDS | LSC | *rps18* |
| **18-nucleotide Repeat** | 37743 | 37790 | 48 | CDS | LSC | *rps18* |
| **24-nucleotide Repeat** | 37699 | 37749 | 51 | CDS | LSC | *rps18* |
| **18-nucleotide Repeat** | 37541 | 37586 | 46 | CDS | LSC | *rps18* |
| **18-nucleotide Repeat** | 37475 | 37539 | 65 | CDS | LSC | *rps18* |
| **18-nucleotide Repeat** | 37433 | 37473 | 41 | CDS | LSC | *rps18* |
| **12-nucleotide Repeat** | 37408 | 37437 | 30 | CDS | LSC | *rps18* |
| **12-nucleotide Repeat** | 37303 | 37332 | 30 | CDS | LSC | *rps18* |
| **12-nucleotide Repeat** | 37210 | 37233 | 24 | CDS | LSC | *rps18* |
| **7-nucleotide Repeat** | 37032 | 37045 | 14 | IGS | LSC | *rps18/rpl33* |
| **9-nucleotide Repeat** | 35784 | 35801 | 18 | IGS | LSC | *petG/trnW-CCA* |
| **8-nucleotide Repeat** | 31181 | 31208 | 28 | IGS | LSC | *trnQ-UUG/psbK* |
| **8-nucleotide Repeat** | 28055 | 28072 | 18 | CDS | LSC | *atpA* |
| **7-nucleotide Repeat** | 25255 | 25273 | 19 | IGS | LSC | *atpF/atpH* |
| **7-nucleotide Repeat** | 12539 | 12552 | 14 | IGS | LSC | *rpoB/trnT-GGU* |
| **7-nucleotide Repeat** | 9203 | 9216 | 14 | IGS/CDS | LSC | *psbC/trnS-UGA* |
| **7-nucleotide Repeat** | 9114 | 9127 | 14 | IGS | LSC | *psbC/trnS-UGA* |
| **8-nucleotide Repeat** | 2059 | 2075 | 17 | CDS/IGS | LSC | *ycf3/psaA* |

**Supplementary Table S7.** Distribution of tandem repeats in the plastome of *C. hildmannianus* subsp. *hildmannianus*. CDS, coding sequences; IGS, intergenic spacers.

| Type | Start | End | Length | Region | Region in the plastome | Location |
| --- | --- | --- | --- | --- | --- | --- |
| 11-nucleotide Repeat | 108826 | 108848 | 23 | intron | SSC | *ndhB* |
| 9-nucleotide Repeat | 103680 | 103698 | 19 | rRNA | SSC | *rrn16* |
| 7-nucleotide Repeat | 102214 | 102227 | 14 | intron | SSC | *trnI-GAU* |
| 8-nucleotide Repeat | 101245 | 101260 | 16 | intron | SSC | *trnA-UGC* |
| 7-nucleotide Repeat | 98749 | 98762 | 14 | rRNA | SSC | *rrn23* |
| 7-nucleotide Repeat | 97119 | 97132 | 14 | IGS | SSC | *trnR-ACG/trnN-GUU* |
| 7-nucleotide Repeat | 92939 | 92952 | 14 | CDS | SSC | *ndhD* |
| 7-nucleotide Repeat | 91017 | 91031 | 15 | IGS | SSC | *ccsA/trnL-UAG* |
| 7-nucleotide Repeat | 88968 | 88981 | 14 | CDS | SSC | *ycf1* |
| 7-nucleotide Repeat | 88500 | 88517 | 18 | CDS | SSC | *ycf1* |
| 24-nucleotide Repeat | 88338 | 88447 | 110 | CDS | SSC | *ycf1* |
| 7-nucleotide Repeat | 88310 | 88388 | 79 | CDS | SSC | *ycf1* |
| 9-nucleotide Repeat | 88259 | 88279 | 21 | CDS | SSC | *ycf1* |
| 9-nucleotide Repeat | 84674 | 84691 | 18 | CDS | IRB | *ycf1* |
| 7-nucleotide Repeat | 84167 | 84181 | 15 | CDS | IRB | *ycf1* |
| 12-nucleotide Repeat | 79197 | 79221 | 25 | CDS | IRB | *ycf2* |
| 9-nucleotide Repeat | 78953 | 78970 | 18 | CDS | IRB | *ycf2* |
| 8-nucleotide Repeat | 76153 | 76168 | 16 | IGS | IRB | *trnH-GUG/psbA* |
| 7-nucleotide Repeat | 71444 | 71457 | 14 | intron | IRB | *rps16* |
| 7-nucleotide Repeat | 71407 | 71420 | 14 | intron | IRB | *rps16* |
| 21-nucleotide Repeat | 69568 | 69609 | 42 | IGS | IRB | *psaI/accD* |
| 19-nucleotide Repeat | 67900 | 67939 | 40 | IGS | IRB | *accD/trnM-CAU* |
| 8-nucleotide Repeat | 61235 | 61250 | 16 | IGS | IRB | *trnE-UUC/trnC-GCA* |
| 9-nucleotide Repeat | 57945 | 57962 | 18 | IGS | IRB | *trnD-GUC/ndhJ* |
| 7-nucleotide Repeat | 52236 | 52251 | 16 | IGS/CDS | LSC | *rpl2/rps19* |
| 21-nucleotide Repeat | 51828 | 51932 | 105 | CDS | LSC | *rps19* |
| 7-nucleotide Repeat | 51810 | 51823 | 14 | IGS/CDS | LSC | *rpl22/rps19* |
| **15-nucleotide Repeat** | 51070 | 51101 | 32 | IGS | LSC | *rpl22/rps3* |
| **7-nucleotide Repeat** | 49061 | 49074 | 14 | IGS | LSC | *rpl26/rpl14* |
| **7-nucleotide Repeat** | 45853 | 45868 | 16 | IGS | LSC | *rpoA/petD* |
| **10-nucleotide Repeat** | 44877 | 44898 | 22 | intron | LSC | *petD* |
| **7-nucleotide Repeat** | 38744 | 38758 | 15 | IGS | LSC | *rps12/rpl20* |
| **8-nucleotide Repeat** | 38139 | 38155 | 17 | IGS | LSC | *rpl20/rps18* |
| **12-nucleotide Repeat** | 37435 | 37464 | 30 | CDS | LSC | *rps18* |
| **12-nucleotide Repeat** | 37342 | 37365 | 24 | CDS | LSC | *rps18* |
| **9-nucleotide Repeat** | 35907 | 35924 | 18 | IGS/CDS | LSC | *trnW-CCA/petG* |
| **8-nucleotide Repeat** | 31306 | 31333 | 28 | IGS | LSC | *trnQ-UUG/psbK* |
| **8-nucleotide Repeat** | 28181 | 28198 | 18 | CDS | LSC | *atpA* |
| **7-nucleotide Repeat** | 25379 | 25397 | 19 | IGS | LSC | *atpF/atpH* |
| **7-nucleotide Repeat** | 12662 | 12675 | 14 | IGS | LSC | *rpoB/trnT-GGU* |
| **7-nucleotide Repeat** | 9329 | 9342 | 14 | IGS/CDS | LSC | *psbC/trnS-UGA* |
| **7-nucleotide Repeat** | 9240 | 9253 | 14 | IGS | LSC | *psbC/trnS-UGA* |
| **8-nucleotide Repeat** | 2191 | 2207 | 17 | CDS/IGS | LSC | *psaA/ycf3* |

## Supplementary Scripts

**Supplementary Script S1.**
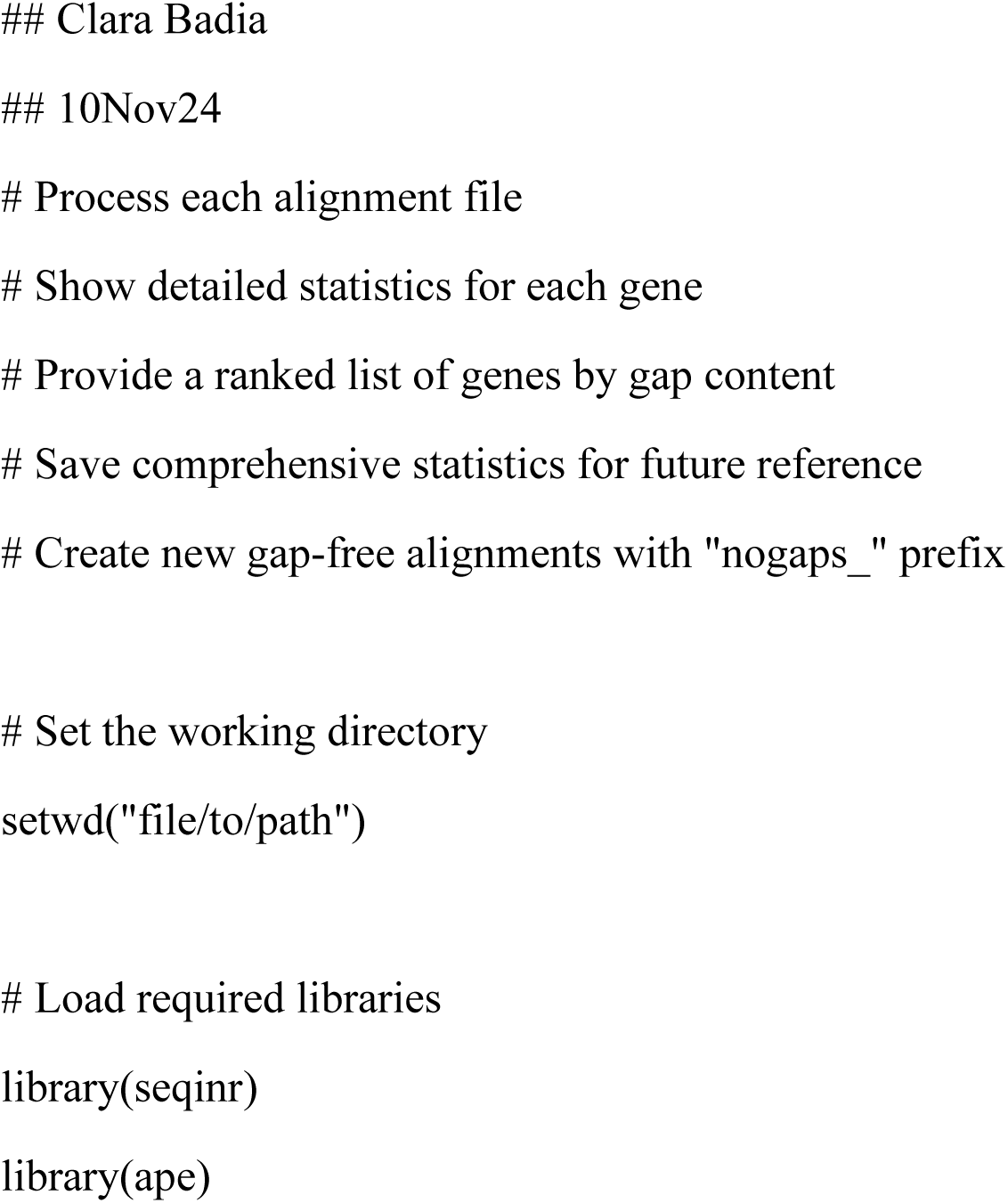
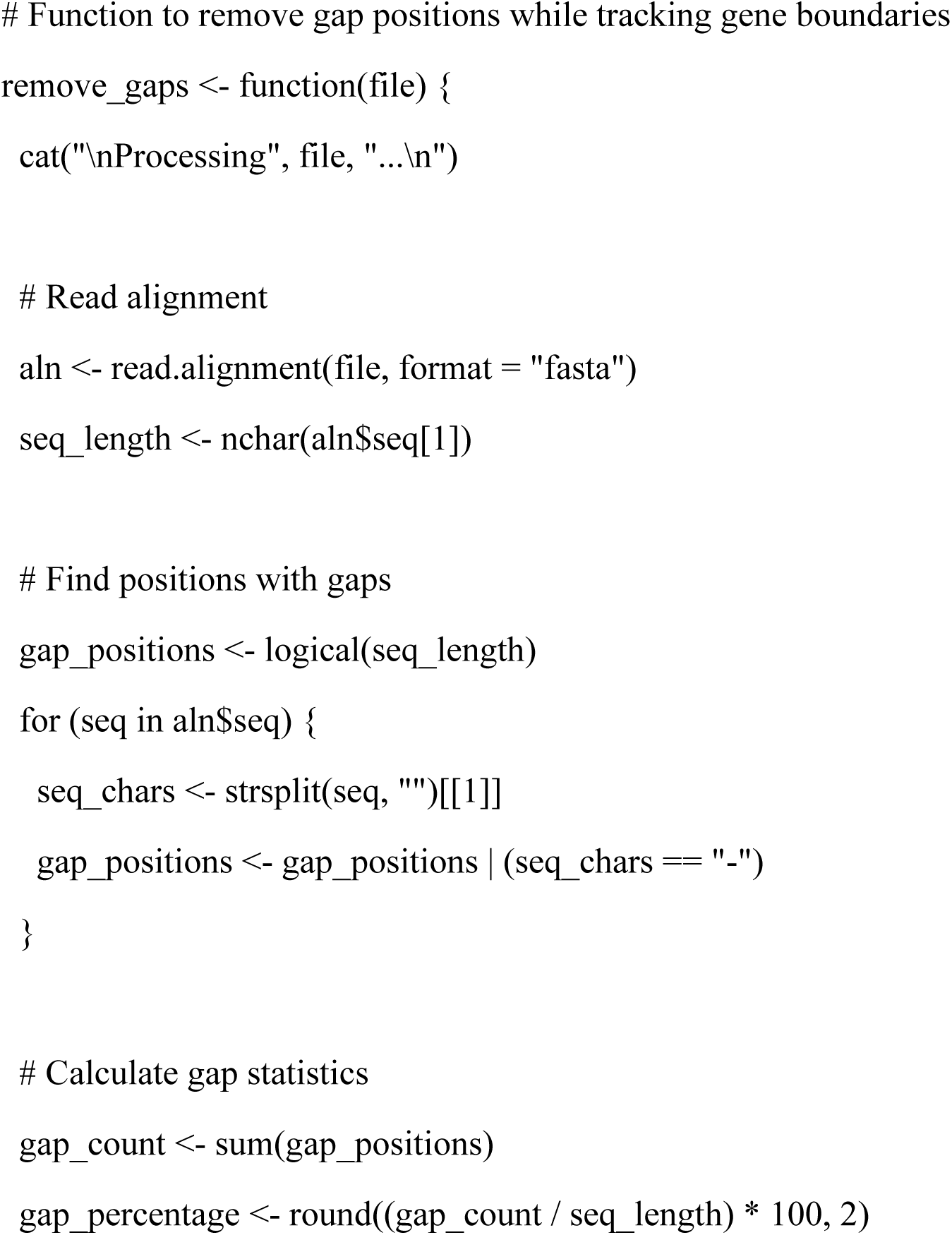
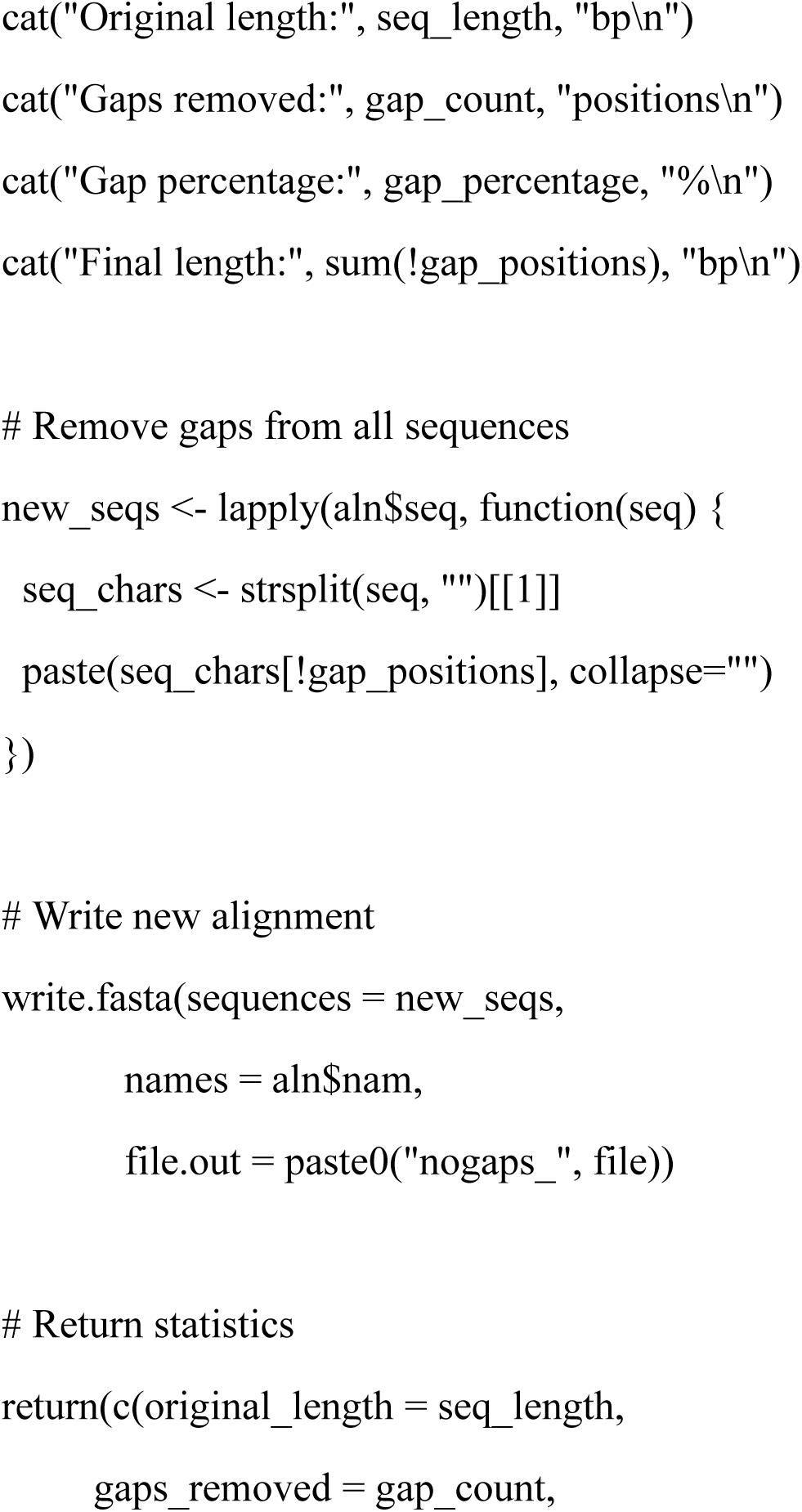
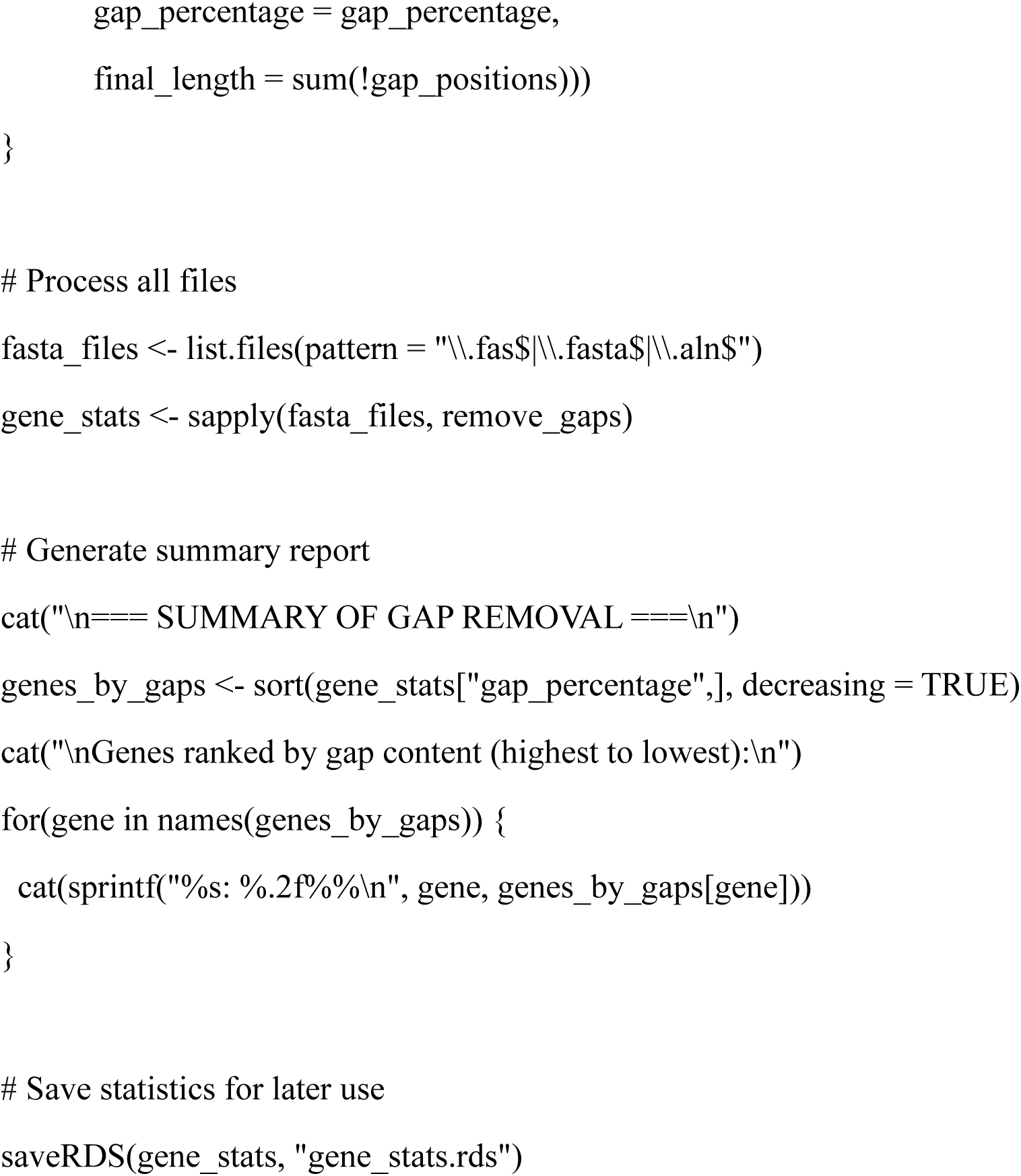
Gap removal script developed in R v. 4.3.3.

